# Evolution of a novel left-right asymmetry in organ size by co-option of a tissue rotation process

**DOI:** 10.1101/2022.01.16.476383

**Authors:** Bénédicte M. Lefèvre, Marine Delvigne, Aurélie Camprodon, Josué Vidal, Virginie Courtier-Orgogozo, Michael Lang

**Author notes:** corresponding authors: Michael Lang, Virginie Courtier-Orgogozo, Bénédicte M. Lefèvre.

## Abstract

Left-right asymmetries have repeatedly evolved in diverse animals and affect the position, shape or size of specific organs. How novel left-right asymmetries arise remains unknown. Here, we examined *Drosophila pachea*, where males have evolved unique asymmetric genitalia lobes and a right-sided copulation posture in the past 3-6 million years. We found that male asymmetric lobes grow in pupae during a 360° clockwise genitalia rotation, a conserved and widespread developmental process in flies. Using two complementary approaches, drug application and a CRISPR-induced MyoID mutant, we altered genitalia rotation and found that asymmetric lobe sizes depend on genitalia rotation completion, while the sidedness of lobe asymmetry is determined by the rotation direction. We then investigated the impact of genital asymmetry on copulation posture. Males with reversed genital asymmetry still mate in the typical right-sided posture, indicating that right-sided behavior is not determined by asymmetric male genitalia. Our study reveals that a novel genitalia asymmetry has evolved through the co-option of a pre-existing tissue remodeling process. Tissue rotation represents a new mechanism, through which a bilateral organ can acquire a left-right size asymmetry.

**Summary statement:** How asymmetric shapes evolve from symmetry is an important question in developmental biology. Using a juvenile hormone derivative, transgenics and CRISPR mutants in the non model species *Drosophila pachea*, we uncover that the developmental process of male genitalia rotation got co-opted during evolution of *D. pachea* to determine the extent and direction of a novel genital left-right asymmetry.

## Introduction

Asymmetries are widespread in the animal kingdom and arise at different biological scales, ranging from molecules to cells, tissues and organs (Blum and Ott, 2018; Palmer, 2009; Schilthuizen, 2013). While the outer body-parts of most bilaterian animals are left-right symmetric, some organs, such as the heart, stomach and spleen in mammals, display left-right asymmetries in position, shape or size (Blum and Ott, 2018; Hamada and Tam, 2020; Palmer, 2009). The default-state development of bilaterians is thought to be symmetric. Left-right asymmetry occurs through symmetry breaking events, involving diverse mechanisms, such as motile cilia and directional fluid flow, orientation of the cell division plane and distribution of molecules within cells (Hamada and Tam, 2020; Lai et al., 2023). Among the different mechanisms involved in left-right asymmetry development, myosin 1 homologs appear to be key factors, both in vertebrates and invertebrates (Hamada and Tam, 2020; Juan et al., 2018; Tingler et al., 2018). In vertebrates such as human, xenopus and zebrafish, myosin 1d homologs are essential for left-right asymmetry establishment (Alsafwani et al., 2021), probably through regulation of the Planar Cell Polarity (PCP) pathway (Juan et al., 2018; Tingler et al., 2018). Myosin 1d in zebrafish appears to regulate motile cilia-driven unidirectional fluid flow and vacuolar trafficking across epithelial cells of the Kupffer’s vesicle, which is a conserved developmental and left-right organizer (Juan et al., 2018; Saydmohammed et al., 2018). In *D. melanogaster*, myosin 1 homologs, in particular Myo1D, have been shown to regulate several left-right asymmetries, either affecting organ position, such as in the Malpighian tubules, or shape, such as the digestive tract and testis coiling (Coutelis et al., 2008). Among the latter, the best characterized are the clockwise looping of the embryonic and adult hindgut (Coutelis et al., 2008; Géminard et al., 2014; Hozumi et al., 2006; Lai et al., 2023; Spéder et al., 2006) and the 360° clockwise rotation of male terminalia during metamorphosis (Spéder et al., 2006; Suzanne et al., 2010). Genitalia rotation is conserved in a wide range of fly species, grouped as cyclorrhapha, to which the *Drosophilidae* family belongs (Feuerborn, 1922; Suzanne et al., 2010). The direction of this rotation is driven by *Myo1D*, encoded by the gene *myo31DF* (Spéder et al., 2006), for simplicity here referred to as *myo1D*. Loss of function of *myo1D* in *D. melanogaster* leads to a reversed, counter-clockwise 360° genitalia rotation (Spéder et al., 2006; Suzanne et al., 2010). Ectopic expression of Myo1D and myosin Myo1C are also sufficient to artificially induce directional twisting of the entire larval body in *D. melanogaster* along the antero-posterior body axis (Lebreton et al., 2018). At cellular scales, *Myo1D* interactions with adherens junctions (Petzoldt et al., 2012; Taniguchi et al., 2011), actin cytoskeleton (Chougule et al., 2020; Lebreton et al., 2018; Petzoldt et al., 2012), planar cell polarity pathway members (González-Morales et al., 2015; Juan et al., 2018; Lai et al., 2023) and Jun kinase pathway regulated apoptosis (Benitez et al., 2010; Macias et al., 2004; Rousset et al., 2010; Taniguchi et al., 2007) have been reported to be important in left-right asymmetry development. Nevertheless, the exact developmental changes underlying the evolution of novel asymmetric morphologies from symmetric organs remains elusive.

Among animals with internal fertilization, genitalia are the most rapidly evolving organs, leading to high interspecific morphological diversity (Eberhard, 2010, 1985; Schilthuizen, 2014). In particular, left-right asymmetric genitalia have evolved independently several times from ancestral symmetric ones (Schilthuizen, 2014, 2013). Genital asymmetry has been hypothesized to evolve in response to changes in mating position (Huber, 2010). Alternatively, asymmetric genitalia may provoke lateralized mating behavior since the asymmetric body parts can mediate the coupling of the female and male abdomen during copulation. However, case-by-case specific assessments are required to test these scenarios. Left-right asymmetric genitalia is rare in *Drosophila*, and has been described in only 11 species among more than 1500 described species (Acurio et al., 2019; Bächli et al., 2021). Among them, a cactophilic species endemic to the Sonoran desert, Mexico, *Drosophila pachea*, displays particularly interesting asymmetries in genitalia. Males have an asymmetric phallus with a right-sided opening for sperm release and a pair of asymmetric external genital lobes, with the left lobe being longer than the right lobe (Acurio et al., 2019; Lang and Orgogozo, 2012; Lefèvre et al., 2021; Pitnick and Heed, 1994). Due to their location at the posterior tip of the male abdomen both at pupal and adult stages, these structures are easy to monitor, which makes them a good model to investigate their development, and in particular, when and how their asymmetry is established. The lobes are not present in the most closely related sister species, *D. nannoptera, D. wassermani*, and *D. acanthoptera*, that diverged 3-6 million years ago from *D. pachea* (Lang et al., 2014), but other asymmetries are found in the sister species. Whereas *D. nannoptera* has fully symmetric genitalia, *D. acanthoptera* possesses an asymmetric phallus and *D. wassermani* has a left-right concave-convex shaped cercus (Acurio et al., 2019; Lang et al., 2014; Pitnick and Heed, 1994; Rice et al., 2019). *Drosophila pachea* mate in an asymmetric, right-sided copulation posture with the male being shifted about 6-8° towards the right side of the female (Acurio et al., 2019; Lang and Orgogozo, 2012; Rhebergen et al., 2016). Surgical manipulations have shown that the lobes increase the chance to establish genital contacts and help to maintain the asymmetric posture throughout mating (Lefèvre et al., 2021; Rhebergen et al., 2016).

In our study, we took advantage of *Drosophila pachea* to investigate how the recently evolved asymmetric male genitalia lobes develop during metamorphosis and monitored lobe growth in fixed tissues and via live-imaging. To investigate whether male genitalia rotation influences asymmetric lobe development, we manipulated male genitalia rotation with application of the juvenile hormone derivative pyriproxyfen. Furthermore, we generated CRISPR-mediated *Myo1D* mutants to invert male genitalia rotation direction and studied rotation *in vivo* in transgenic *D. pachea* expressing a fluorescent membrane marker. Finally, we investigated the effect of reverse left-right genitalia anatomy on copulation posture.

## Results

### The asymmetric genital lobes of *D. pachea* males form during genitalia rotation

*D. pachea* adult males exhibit asymmetric lobes (Figure 1A), with the left lobe being about 1.5-fold longer than the right lobe (Lang and Orgogozo, 2012; Lefèvre et al., 2021; Rhebergen et al., 2016) (Figure 1A). This size asymmetry can be due to various differences in the development of the left and right lobes, related to cell division rates, cell death rates, cell sizes, timing of cell proliferation, or cell recruitment or intercalation from neighboring tissues. To characterize the growth of these lobes, we performed immunostainings of male genitalia at various time points during pupal development, from 24 h to 48 h after puparium formation (APF, Figure 1B, Supplementary Dataset 1). Apoptotic cells were identified by immunostaining using a cleaved caspase 3 specific antibody (Figure S1), and were sporadically found in the lobe tissues at 28 h APF, 31 h APF and at 48 h APF, suggesting that differential apoptosis between left and right lobes does not contribute to their asymmetry. The number of nuclei and mitotic cells were quantified in each lobe by automatic detection of nuclei in manually-segmented lobe volumes (Figures 1C and D). We observed that the lobes form as a fold at the junction of the so called hypandrial sheet that covers the internal parts of the genitalia, the dorsally located surstili and the ventral epandrial arch, also called lateral plate. We defined the proximal limits of the lobes as the line delimiting the lateral border of the surstili (Figure 1B, green line). Lobes can be detected as small, rudimentary buds at 24 h APF. At later stages, lobe tissue was observed to increase in cell number on both sides, and a pronounced difference in cell number between the left and right lobes was detectable at 36 h APF, which was roughly maintained constant until 48 h APF with approximately 400 cells in the left lobe and 200 cells in the right one (Figure 1C). The shape of the left lobe further differentiated between 43 h APF and 48 h APF to adopt a curved z-shape, not observed in the right lobe (Figure 1B). Mean lobe cell size was calculated for each lobe at different stages as the ratio of estimated lobe volume and the number of nuclei counted therein (n = 46, Figure S1 A,B). Cell sizes were only slightly higher in the left lobe at 124 µm^3^ than in the right lobe at 106 µm^3^ (t-test, t = 2.2973, df = 75.736, p = 0.02436) and may contribute only slightly to the observed lobe size difference.

**Figure 1:**
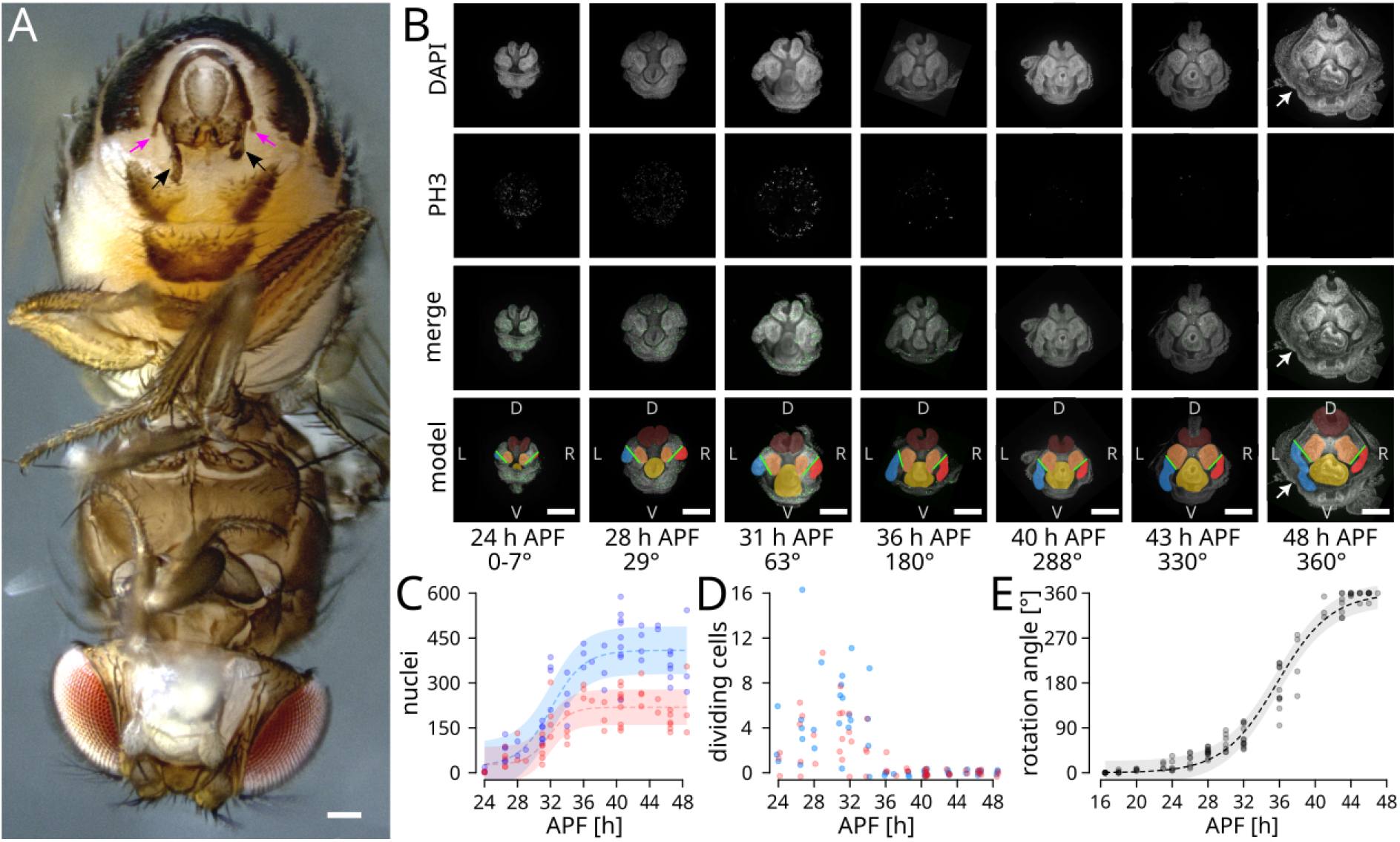
Asymmetric *D. pachea* male genital lobes grow during clockwise genitalia rotation. **(A)** Wild-type *D. pachea* adult male in ventral view with external male genitalia on the top. Asymmetric lobes are indicated with black arrows, lateral spines with magenta colored arrows. **(B)** Lobe formation in *D. pachea* wild-type stock 1698.01 between 24 h after puparium formation (APF) and 48 h APF. Nuclei were stained with DAPI (grey, first and third rows) and mitotic cells with anti-phospho-histone H3 antibody (grey in second row, green in third row). Genital primordial structures are highlighted (last row): anal plates (brown), surstili (orange), phallus (yellow), left lobe (blue) and right lobe (red). The proximal limits of the lobes (green lines) are defined as the ventral - lateral border of the surstili. The white arrows (first, third and last row) indicate the z-shaped left lobe at 48 h APF. Body axes are indicated with L = left, R = right, D = dorsal, V = ventral. The average clockwise genitalia rotation angle is indicated below each column. The scale bar is 100 μm. **(C)** Number of nuclei in the left (blue) and right (red) lobe at different developmental time points (Figure 1B). 4-parameter logistic regressions and standard errors are shown as dashed lines and colored shades in blue and red, respectively. **(D)** Number of phospho-histone H3-positive cells in the left (light blue) and right (pink) lobe during pupal development. **(E)** Genitalia rotation over time in *D. pachea* male wild-type stock 1698.01. Genitalia orientation is the angle of the main axis of the surstili relative to the main axis of the developing legs. A 2-parameter logistic regression curve is shown as a dashed line, the gray shade is the standard error.

Our observations indicate that both the left and right lobes grow during the same time period. Cell proliferation was primarily observed between 24 h APF and 31 h APF and was progressively decreased at later time points, to become non detectable at 40 h APF and later time points (Figure 1D). We observed no significant difference between lobes in the proportion of dividing cells with respect to the total numbers of cells. Although we cannot exclude the existence of a short bout of proliferation that we may have missed and that would vary between left and right sides, our data suggests that the final difference in cell number between lobes is not due to proliferation but to other mechanisms such as differential cell intercalation from neighboring tissues.

The clockwise rotation of male genitalia occurs during pupal development and is a widespread phenomenon in flies (Feuerborn, 1922; Petzoldt et al., 2012; Spéder et al., 2006; Suzanne et al., 2010). To investigate a potential link between genitalia rotation and asymmetric lobe growth, we first wanted to check if lobe growth occurs during, before, or after genitalia rotation. We monitored the rotation of genitalia in *D. pachea* on dissected posterior parts of male pupae at various time points during pupal development (Supplementary Dataset 2). In particular, we approximated the dorso-ventral axis of male genitalia as the midline between the dorsal anal plates and the ventral surstyli (Figure S2 A). Then, we measured the orientation of this axis relative to the dorso-ventral body midline relative to the ventrally located leg primordia (Figure S2 A). We found that *D. pachea* genitalia rotation occurs between 20 h APF and 45 h APF, with 180° half-rotation at approximately 36 h APF. Our results reveal that male genital lobes in *D. pachea* grow exactly during genitalia rotation, and that lobe asymmetry becomes progressively apparent during the first half-rotation.

### Inhibiting genitalia rotation with the juvenile hormone derivative pyriproxyfen alters genitalia rotation and adult lobe size

Genitalia rotation is widespread in flies but only *D. pachea* possesses asymmetric genitalia lobes. We hypothesized that this rotation process may have been co-opted during evolution of *D. pachea* to trigger asymmetric tissue growth in the male genital lobes. We therefore sought to perturb genital rotation, using both chemical and genetic means, and examine the effects on development of the genital lobe size and asymmetry. First, we conducted a drug treatment that could perturb genitalia rotation (Supplementary Dataset 3). Juvenile hormone derivatives such as pyriproxyfen are used as insecticides and were shown to affect metamorphosis and to partially inhibit male genitalia rotation in *D. melanogaster* (Ádám et al., 2003; Riddiford and Ashburner, 1991). First, we conducted drug application on the dorsal pupal surface as previously reported (Ádám et al., 2003; Riddiford and Ashburner, 1991) and evaluated survival (Figure 2A) and orientation of adult male genitalia (Figure 2B). Similar to previous measurements to track male genitalia rotation in pupae, we approximated the dorso-ventral axis of male genitalia as the midline between the dorsal anal plates and the ventral surstyli (Figure S2 B,C) but now calculated the orientation of this axis relative to the antero-posterior adult body midline. The estimated half lethal dose (LD50) to pyriproxyfen of *D. pachea* was 0.123 pmol, an order of magnitude lower than in *D. melanogaster* with a LD50 of 1.467 pmol, and no perturbation of genitalia rotation could be detected (Figure 2A). To limit mortality and observe an effect of pyriproxyfen, we then tried to restrict application of pyriproxyfen to the posterior tip of *D. pachea* pupae. This locally concentrated the drug in close proximity to genitalia and had effects on their development, although a slightly lower LD50 value of 0.061 pmol was found. While genitalia orientation appeared to be normal when applying solvent alone (Figure 2B, n = 16), upon treatment with 0.06 to 0.07 pmol pyriproxyfen, genitalia rotated partially, leading to adult genitalia with orientation deviating from 0° to 235° compared to the wild-type orientation (Figure 2B-E, Supplementary Dataset 4).

**Figure 2:**
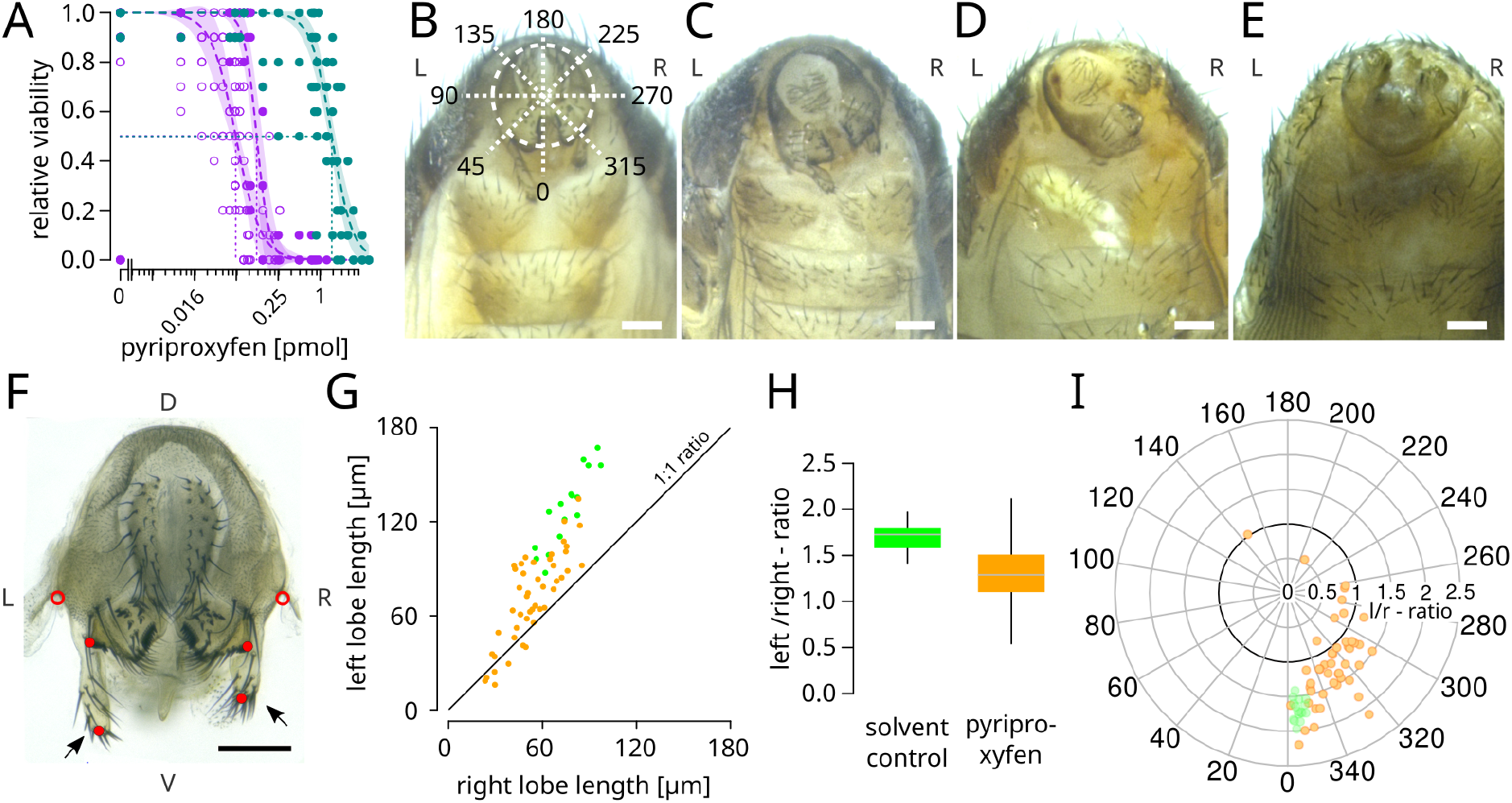
The lobe length ratio increases with male genitalia rotation progress. **(F)** Viability half lethal dose (LD50) in pyriproxyfen treatment with dorsal application, *D. melanogaster* (cyan) 1.467 pmol and of *D. pachea* (purple) 0.123 pmol. *D. pachea* viability to applications to the the posterior tip of the pupae (purple, open circles) was 0.061 pmol. Logistic regression curves and confidence intervals are shown. **(B-E)** Abdomen of *D. pachea* adult males in ventral view, treated with (B) solvent, or (C-E) 0.6 - 0.7 pmol pyriproxyfen dissolved in acetone. The polar diagram in (B) indicates genitalia orientation coordinates. Left and right are indicated with L and R, respectively **(F)** Dissected *D. pachea* male genitalia. The lobes are indicated by arrows. Measurement landmarks are indicated by red dots. Lobe length is measured as the distance between the tip of each lobe and the basal lateral edge of each surstylus. In other studies, lobe length was measured from the tip of the lobe to the base of the lateral epandrial spines, indicated by the red open circle. Body axes are indicated with L = left, R = right, D = dorsal, V = ventral. **(G)** Scatter plot of left and right lobe length in pyriproxyfen treated individuals (orange) and solvent control individuals (green). Left and right lobe length are highly correlated (pearson correlation coefficient 0.90922, p < 0.0000001). **(H)** Left-right lobe ratio in pyriproxyfen treated individuals (orange) and solvent control individuals (green) of data points shown in (F). **(I)** Left-right lobe length ratio and genitalia orientation angle in pyriproxyfen treated individuals (orange) and solvent control individuals (green) presented on a polar diagram. The lobe length ratio is correlated with genitalia orientation (Pearson correlation, coefficient 0.53295, p< 0.00001). The bold line corresponds to the 1:1 length ratio. The scale bar is 100 µm.

We carried out length measurements of both lobes (Figure 2F, Supplementary Dataset 4). In previous studies (Lang and Orgogozo, 2012; Lefèvre et al., 2021; Rhebergen et al., 2016), lobe length was measured as the distance from the distal tip of the lobe to the base of a lateral genital spine (Figure 2E). However, spines were not observed in most treated individuals, so we used the surstili as an alternative and common reference point to define the proximal end of the lobe (Figure 2E). Among pyriproxyfen-treated males with genitalia rotation phenotypes, left and right lobe length variation was strongly correlated (Figure 2G). More precisely, the more genitalia orientation deviated from normal orientations, the shorter both lobes were, indicating that growth of both lobes was generally inhibited to various extents by the treatment. Overall, solvent control treated individuals had a lobe length ratio of 1.70 ± 0.148 (mean ± standard deviation, n = 16) (Figure 2 H), while pyriproxyfen treated males revealed a lower length ratio of 1.32 ± 0.364 (median ± standard deviation, n = 46). In addition, the ratio of left and right lobe length was correlated with genitalia orientation (Figure 2 I), compared to control treatment with solvent application. Altogether, our data show that the juvenile hormone derivative pyriproxyfen blocks both genitalia lobe development and asymmetric tissue growth.

### Inverting genitalia rotation direction via *Myo1D* mutation reverts left-right asymmetry in *D. pachea* male genitalia

One possibility is that pyriproxyfen blocks genitalia lobe development and that asymmetric lobe growth is dependent upon the completion of genitalia rotation. However, pyriproxyfen drug treatment can also affect other processes besides genitalia rotation, so it is not clear if the defects we observed in lobe size are due to the alteration of genitalia rotation or to other effects of the pyriproxyfen treatment. For example, anal plates (cerci) were observed to be less differentiated in pyriproxifen-treated males than in controls (Figure 2E), suggesting that pyriproxifen may hamper various developmental processes and not just genitalia rotation. To further explore the role of genitalia rotation on asymmetric lobe growth, we aimed to manipulate the rotation direction by another means, genetically. Genitalia rotation has been characterized in *D. melanogaster*, and its clockwise direction is driven by Myo1D, as loss of function of the encoding gene leads to counter-clockwise rotation (Petzoldt et al., 2012; Spéder et al., 2006; Suzanne et al., 2010). We hypothesized that mutation of *myo1D* in *D. pachea* would lead to counter-clockwise male genitalia rotation as in *D. melanogaster*. We thus generated a CRISPR/Cas9-mediated mutation of *D. pachea myo1D* (Bassett and Liu, 2014; Jinek et al., 2012) (*myo1D*^*mut*^ allele). Sequencing revealed a 12-bp deletion and a 7-bp insertion in the seventh coding exon, resulting in a frameshift and a premature stop codon (Materials and Methods, Figure S3). Heterozygous *myo1D*^*wt/mut*^ males did not have visible alterations of male genitalia morphology and orientation (n = 69) and were indistinguishable from *myo1D*^*wt/wt*^ males (n = 41) (Supplementary Dataset 4). In contrast, homozygous *myo1D*^*mut/mut*^ males (n = 85) showed aberrant genitalia orientation phenotypes (Figure 3A-D) and the asymmetry direction of the left and right genital lobe was reversed, with a longer right lobe and a shorter left lobe (Figure 3E). Left and right lobe lengths were strongly correlated in *myo1D*^*wt/mut*^ and *myo1D*^*wt/wt*^ males (Figure 3F), but only weakly in *myo1D*^*mut/mut*^ males. Overall, *myo1D*^*wt/wt*^ and *myo1D*^*wt/mut*^ males had similar lobe length ratios of 1.87 ± 0.177 (mean ± standard deviation, n = 41) and 1.90 ± 0.205 (median ± standard deviation, n = 70) (Figure 3G), respectively, while *Myo1D*^*mut/mut*^ males revealed an inverted length ratio of 0.57 ± 0.149 (median ± standard deviation, n = 87). The ratio of left and right lobe length was not significantly correlated with genitalia orientation in *myo1D*^*mut/mut*^ males (Figure 3H, Figure S4). While right lobe length was indeed correlated with the genitalia orientation angle, left lobe length appeared to vary independently so that the effect on the length ratio was probably too weak to be detectable (Figure S4). In one *myo1D*^*mut/mut*^ male, the genital lobe asymmetry was not reversed: although lobe lengths were similar, genitalia were mis-oriented and the lobes were strongly bent. It was unclear if genitalia rotated clockwise or counter-clockwise during development of this individual. Remnant activity and expression of the *myo1D*^*mut*^ allele, could have potentially led to partial rotation in clockwise direction.

**Figure 3:**
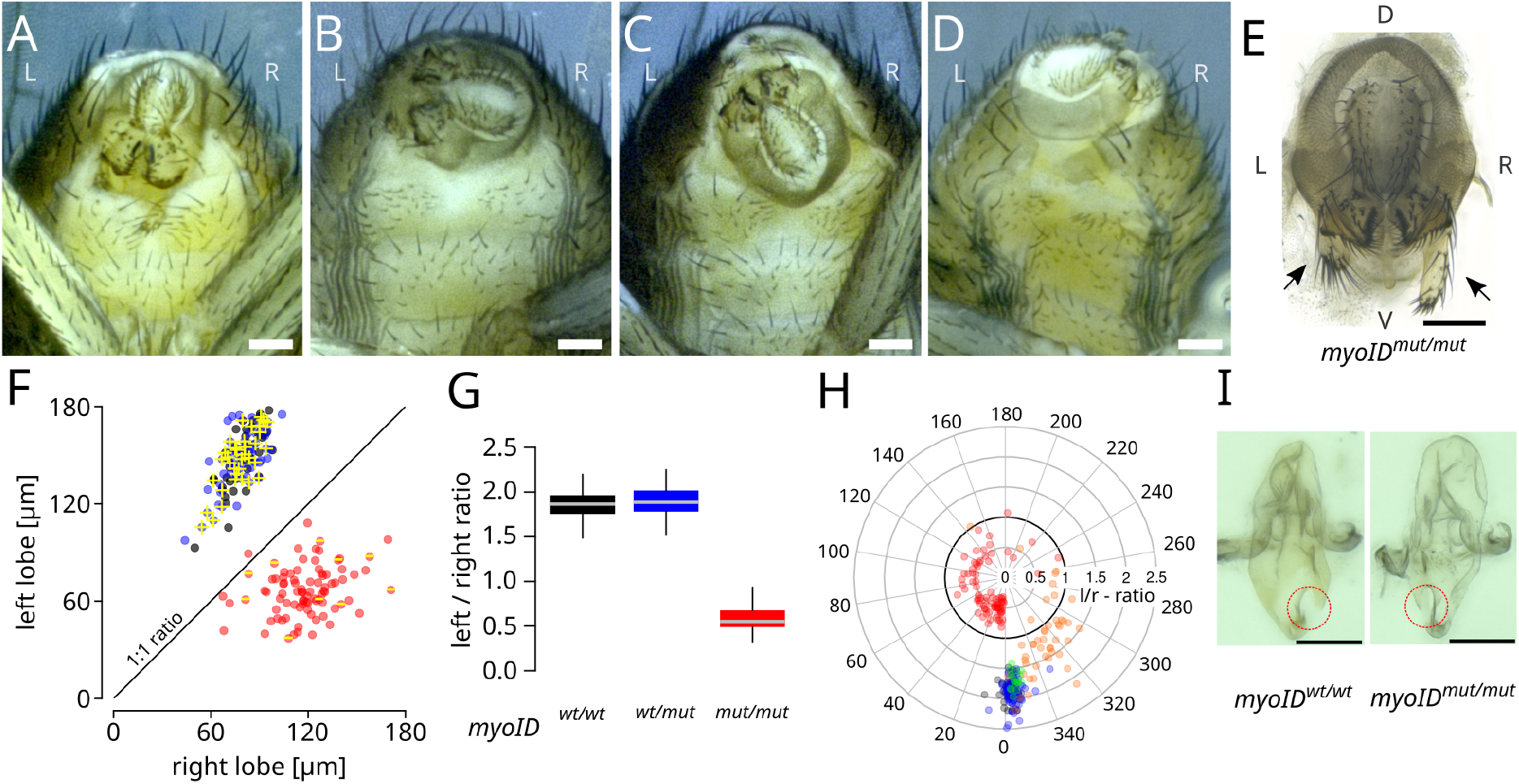
*D. pachea Myo1D* mutant males have reversed left-right genitalia. **(A-D)** Abdomen of *D. pachea myo*^*mut/mut*^ males in ventral view. Left and right are indicated with L and R, respectively. **(E)** Dissected *myo*^*mut/mut*^ male genitalia with reverted lobe asymmetry. Lobes are indicated by arrows. Body axes are indicated with L = left, R = right, D = dorsal, V = ventral. **(F)** Scatter plot of left and right lobe lengths of *myo*^*wt/wt*^ males (black), *myo*^*wt/mut*^ males (blue), and *myo*^*mut/mut*^ males (red). Left and right lobe length are highly correlated in *myo*^*wt/wt*^ and *myo*^*wt/mut*^ males (pearson correlation coefficient 0.711, p < 0.0000001) but weakly correlated in *myo*^*mut/mut*^ males (pearson correlation coefficient 0.256, p < 0.017). Genitalia rotation direction (‘+’ = clockwise, ‘-’ = counter-clockwise) is annotated for some individuals in yellow. **(G)** Left-right lobe length ratio of *myo*^*wt/wt*^ males (black), *myo*^*wt/mut*^ males (blue), and *myo*^*mut/mut*^ males (red) of data points shown in (F). **(H)** Left-right lobe length ratio and genitalia orientation angle of *myo*^*wt/wt*^ males (black), pyriproxyfen treated individuals (orange), solvent control individuals (green), *myo*^*wt/mut*^ males (blue), and *myo*^*mut/mut*^ males (red) presented on a polar diagram. The bold line corresponds to the 1:1 length ratio. The scale bar is 100 µm. **(I)** Dissected *D. pachea* phalli of a *myo*^*wt/wt*^ male and a *myo*^*mut/mut*^ male. The opening for sperm release (phallotrema) is indicated with a red circle.

To monitor genitalia rotation direction *in vivo*, we generated and integrated a DE-Cadherin::EYFP fusion construct into *D. pachea* (DE-Cad::EYFP_2) (Material and Methods, Figure S5). The DE-Cad::EYFP_2 was placed in a piggybac vector (Horn et al., 2003) and was inserted at a random genome position into the *myo1D*^*mut/mut*^ stock by germline transformation (Figure S5). We examined male genitalia rotation in pupae by time-lapse imaging (Supplementary Dataset 5). We used heterozygous DE-Cad::EYFP individuals to reduce potentially deleterious effects of the piggybac insert (Figure S6). All *myo1D*^*wt/wt*^ (n = 9) and *myo1D*^*wt/mut*^ (n = 25) males underwent clockwise and complete genitalia rotation and had a wild-type lobe length ratio (Figure 3G, Supplementary Dataset 4). In contrast, genitalia rotation progress was variable for *myo1D*^*mut/mut*^ males (Supplementary Dataset 4) and the rotation direction was counter-clockwise in all monitored individuals (n = 9), Figure 3F). Therefore, in *D. pachea* Myo1D controls the direction of genitalia rotation as in *D. melanogaster*. Rotation direction is associated with the lobe asymmetry ratio and Myo1D appears to determine the direction of left-right asymmetry in male genitalia indirectly through its action on the direction of genitalia rotation.

The phallus asymmetry was also reversed in *myo1D*^*mut/mut*^ males. The opening for sperm release (phallotrema) is consistently located at the right distal tip of the phallus (Acurio et al., 2019) in *myo1D*^*wt/wt*^ males (n = 9) and *myo1D*^*wt/mut*^ (n = 13), while it is located at the left tip of the phallus in *myo1D*^*mut/mut*^ males (n = 29) (Figure 3I) (Supplementary Dataset 4). Summarizing, these data indicate that *Myo1D*-mediated regulation of male genitalia rotation is conserved between *D. melanogaster* and *D. pachea*. In addition, perturbation of genitalia rotation direction through a *myo1D* loss of function mutation is sufficient to reverse genitalia asymmetry both in the lobes and in the phallus.

### Direction of asymmetric male genitalia and asymmetric mating posture are uncoupled in *D. pachea*

Asymmetric genitalia evolved independently several times in animals (Schilthuizen, 2014, 2013). In insects, it has been proposed that genitalia evolve due to changes in mating posture (Huber, 2010). Evolution of both, mating posture and genitalia are subject to sexual selection, but the concrete interactions driving their possible co-evolution are not well understood. In addition to asymmetric male genitalia, *D. pachea* also adopts a right-sided mating posture in which the male is shifted 6-8° to the right side of the female body midline (Acurio et al., 2019; Lang and Orgogozo, 2012; Lefèvre et al., 2021; Rhebergen et al., 2016) (Figure 4A). We aimed to investigate the effect of reversed asymmetric male genitalia on the lateralized mating posture. For this, we monitored courtship and copulation of *myo1D*^*wt/wt*^ and *myo1D*^*wt/mut*^ males with wild-type genital asymmetry, and *myo1D*^*mut/mut*^ males with reversed genitalia asymmetry. We placed single males together with a *myo1D*^*wt/wt*^ female and monitored behavior during courtship for at most 60 min or until copulation ended (Supplementary Dataset 7, Figure S7). Most monitored couples with *myo1D*^*wt/wt*^ (n = 11/12) and *myo1D*^*wt/mut*^ (n = 13/13) males achieved a stable copulation posture within this period, in contrast to only 31% of couples with *myo1D*^*mut/mut*^ males (n = 10/32) (Figure S7). Among the *myo1D*^*mut/mut*^ males which copulated, no major genitalia orientation difference was detected compared to *myo1D*^*wt/wt*^ males, but orientations significantly deviated in males that failed to copulate (Figure S7B).

**Figure 4:**
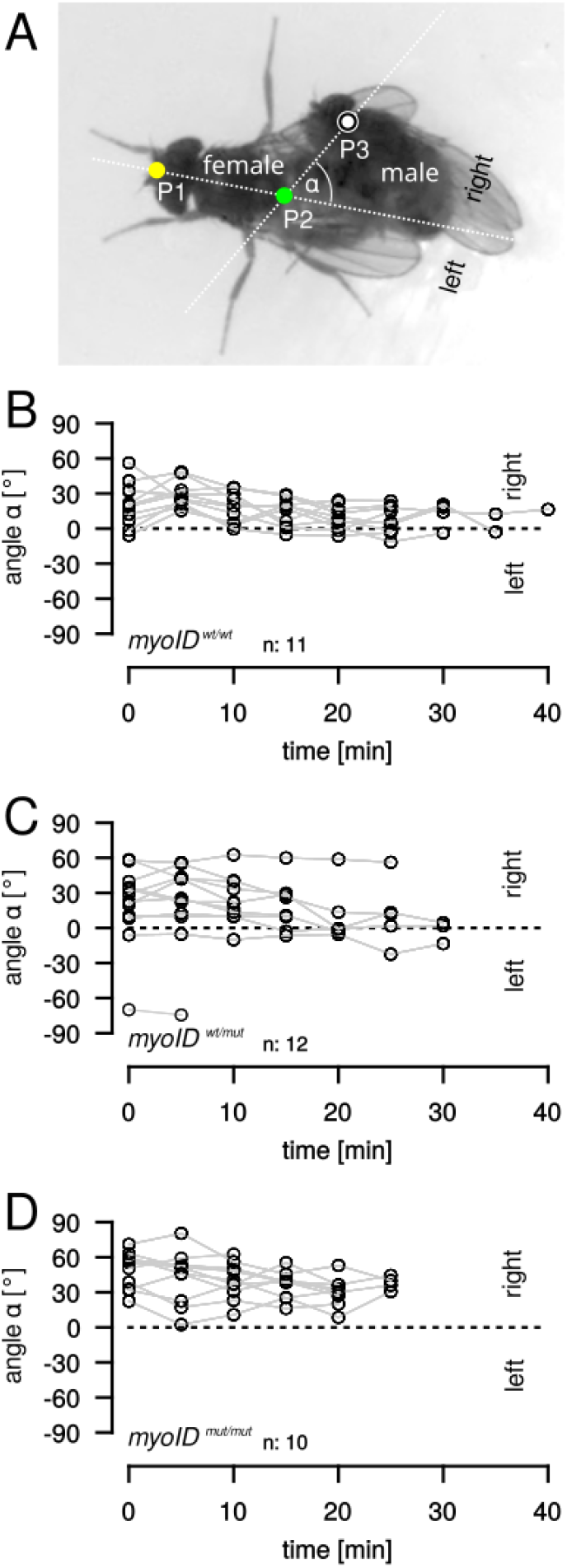
The right-sided *D. pachea* mating posture does not depend on the direction of male genital asymmetry. (A) Copulating *D. pachea* couple. The position of the male relative to the female midline was estimated on extracted images of movies, as in (Acurio et al. 2019), based on three landmark positions on the female and male body: P1 (yellow point) indicates the anterior medial tip of the female head, P2 (green point) the distal tip of the female scutellum and P3 (white open circle) corresponds to the most posterior medial point of the male head. The angle (α) between white lines (P1-P2) and (P2-P3) was used to measure sidedness of the mating position, positive values: right-sided, negative values: left-sided. **(B-D)** Mating angles across copulation of wildtype females with males: *myo1D*^*wt/wt*^ with the wild-type lobe length ratio (B), *myo1D*^*wt/mut*^ heterozygous males with wildtype lobe length ratio, (C) *myo1D*^*mut/mut*^ males with reverted lobe asymmetry ratio. The hypothesis angle = 0 was rejected for each male genotype (GLM fit angle ∼ genotype, all p < 1 × 10^−10^). Positive angle values correspond to right-sided orientations of the male head relative to the female. Data points that correspond to the same trial are connected by gray lines. The number of analyzed mating couples is indicated by n.

We examined the position of the male head relative to the female antero-posterior midline at regular time points during copulation (Supplementary Dataset 8, Figure 4, Figure S8). Regardless of the genotype and direction of genital asymmetry, *myo1D*^*wt/wt*^, *myo1D*^*wt/mut*^ and *myo1D*^*mut/mut*^ males consistently adopted a right-sided copulation posture (Figure 4). This result reveals that right-sided mating posture in *D. pachea* does not depend on the direction of male genital asymmetry. Direction of male asymmetric genitalia and sidedness of mating posture are thus uncoupled in *D. pachea*.

## Discussion

Working on non-model species to investigate animal development has become easier in the last decade with the advance in genetic engineering (Courtier-Orgogozo and Martin, 2020). It opens new opportunities to explore questions for which the classical model species are not the most well suited. *Drosophila pachea* constitutes an excellent emerging model to study left-right asymmetry with its asymmetric genitalia and asymmetric mating posture (Acurio et al., 2019; Lang and Orgogozo, 2012; Lefèvre et al., 2021; Rhebergen et al., 2016), two traits that are absent in the model species *D. melanogaster*. It is still challenging to raise and manipulate diverse *Drosophila* species with non-conventional food requirements and to establish and maintain transgenic lines without chromosomal balancers, although new protocols have become available to overcome the latter (Stern, 2022). Effort is ongoing to further adapt and simplify genetic tools, originally designed for model-species, to become applicable for *D. pachea* and other species of choice.

We wanted to explore how a new left-right asymmetry arises during evolution. We observed that two asymmetric developmental processes, asymmetric genital lobe growth and genitalia rotation, occur at the same time. We thus investigated if the conserved genitalia rotation may have been co-opted during evolution of *D. pachea* to direct the novel left-right asymmetry in male genital lobes. Irrespective of the approach we used to perturb genitalia rotation, either pyriproxyfen treatment or *myo1D* mutants, we found that asymmetric lobe growth is dependent upon rotation completion, and that asymmetry direction of genitalia is imposed by genitalia rotation direction. Changes in mechanical cell-cell or tissue interactions, or in the timing of their interactions, may be cues for morphological diversification. The rotation direction can potentially be translated into morphological left-right asymmetry through direct mechanical interactions between the rotating and non-rotating tissues, via frictional resistance between rotating and non-rotating tissues (mechanical model, Figure 5A). Since the growth direction of the lobes is towards the ventral side of the genitalia, the clockwise rotation may induce pulling forces that are aligned with the growth direction of the left lobe and enhance its growth, but that are in opposite direction for the right lobe and might thus limit its growth. Indeed, live-imaging of rotating genitalia reveals an intermediate accumulation of a tissue bud in movies with *myo1D*^*wt/mut*^ and *myo1D*^*wt/wt*^ males (n = 34), in the close surroundings of rotating genitalia primordia (Figure S9, movie 1), approximately from 90° to 180° of wild-type rotation and when lobe asymmetry becomes apparent. In *myo1D*^*mut/mut*^ males (n=9), a similar bud is observed in a similar orientation (Figure S10, movie 2) but appears during the second half of rotation, between 180° and 270°. Global tissue interactions of different parts of genitalia can affect genitalia development. One known case has been reported for *D. melanogaster* and closely related species that share another unique male genitalia structure called posterior lobes, located centrally between the surstili and the dorsal cerci (anal plates). Interspecific shape variation of the posterior lobes was reported to depend on changed mechanical interactions of genital tissue and the covering extracellular matrix, inducing a tinkering of cell size and shapes (Smith et al., 2020). Here, we propose that lobe asymmetry results from left-right differential force transmission in the genitalia, due to interactions between rotating and non rotating parts. Alternatively, rotation direction may translate a transient dorso-ventral gradient of signaling factors into left-right information (signaling model, Figure 5B). For example, one can imagine a dorsally-located growth promoting factor that is more concentrated or more active during the first half of the rotation process. The rotation will bring the left lobe into contact with this growth promoting factor during the first half of the rotation and then the right lobe during the second half of rotation, when the activating factor is at lower concentrations. Alternatively, one can imagine a ventrally-located growth inhibitory factor that is active during the first half of the rotation process, or a dorsally-located growth inhibitory factor active during the second half. Under this model, changing the direction of rotation can reverse asymmetry by switching the timing of exposition between the left and right sides. One way or another, evolution of this novel morphological asymmetry in lobes appears to be the consequence of complex tissue interactions and remodeling of the entire genital primordium.

**Figure 5:**
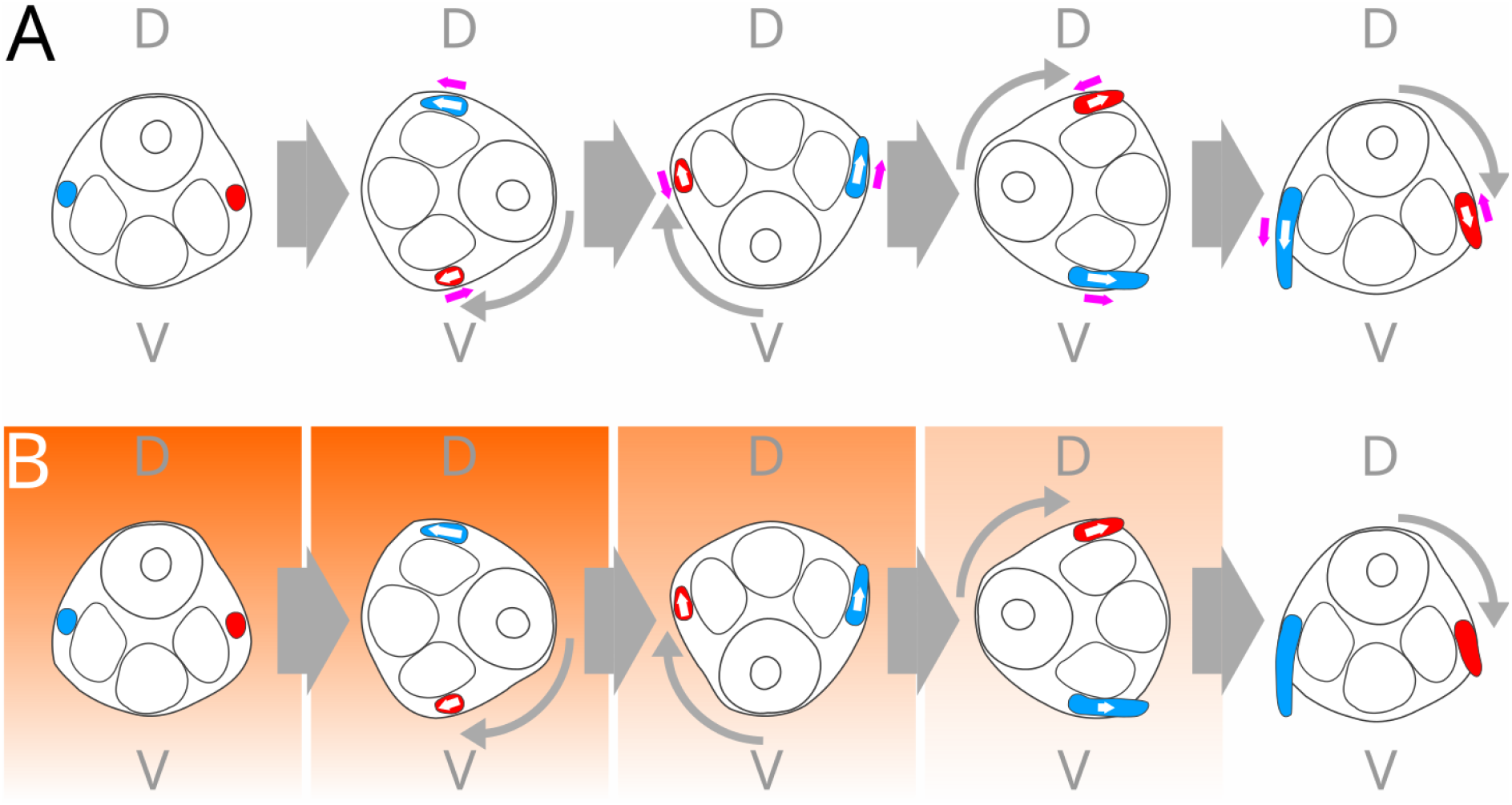
Two models on how the direction of genitalia rotation may determine the direction of asymmetry in *D. pachea* male genitalia lobes. Male genitalia during metamorphosis are outlined, the left lobe is highlighted in blue and the right lobe in red, D: dorsal, V: ventral, narrow gray arrows indicate genitalia rotation direction, bold arrows time. Directions of lobe growth are indicated with white arrows. **(A)** Model 1: Friction force is assumed to act on the lobes (directions indicated by magenta arrows) due to physical interaction of rotating and non-rotating tissues. The orientation of friction force is similar to the growth direction of the left lobe but opposite to the direction of right lobe growth, due to the clockwise rotation direction. This favors tissue growth in the left lobe while it inhibits growth in the right lobe (indicated by the length of the magenta arrows). **(B)** Model 2: A growth factor (orange) is hypothesized to accumulate transiently in the dorsal region of the animal, but decreases with time. Due to clockwise-rotation, the left lobe comes into contact with the growth factor first and enhances growth. When the right lobe enters the dorsal region, the growth factor is no longer present or less concentrated, leading to a right lobe smaller than the left lobe. Differential local concentrations of this signaling molecule lead to enhanced growth (indicated by the length of white arrows), first at the left lobe, then at lower levels at the right lobe due to reduced levels of the signaling factor.

This study provides the first experimental insights into the developmental origin of a recently evolved left-right asymmetry, involving the recruitment of a conserved, pre-existing asymmetric rotational developmental process to derive a new, left-right asymmetric structure. Distinct genital asymmetries are also found in two of three closely related sister species of *D. pachea*: *D. acanthoptera* has an asymmetric phallus (Acurio et al., 2019; Vilela and Bächli, 1990), *D. wassermani* has concave convex shaped anal plates (Lang et al., 2014; Pitnick and Heed, 1994). In contrast, no asymmetries were detected in the male genitalia of *D. nannoptera* and in the more distanly related species *D. bromeliae* and *D. machalilla* (Acurio et al., 2019). Recent phylogenies (Lang et al., 2014; Suvorov et al., 2022) are also consistent with a single evolutionary transition from symmetric to asymmetric genitalia in the common ancestor of *D. pachea, D. acanthoptera* and *D. wassermani*. Genitalia rotation can be a common and conserved developmental cue that may affect distinct developmental processes in each species that have later diversified during their recent, lineage specific evolution. To test this, it would be interesting to create *myo1D* mutants in *D. acanthoptera* and *D. wassermani*.

An intriguing point is that, while genitalia rotation is widespread in Cyclorrhaphan flies (Suzanne et al., 2010), it does not lead to genital asymmetry in most species (Huber et al., 2007). For example, only eight cases of male genital asymmetry are known among more than 1500 described *Drosophila* species (Acurio et al., 2019; Bächli et al., 2021). It means that in most cases, the clockwise genitalia rotation does not lead to asymmetric development of left and right sides and that development of male genitalia symmetry is generally robust to signaling or mechanical constraints, with just a few exceptions. Under the mechanical model (Figure 5A), physical constraints act between rotating genitalia and non-rotating outer tissues. A mediating component may connect rotating and non-rotating tissues only in species that develop genital asymmetry. This could possibly be mediated by extracellular matrix components. As mentioned above, the male genital primordium of *D. melanogaster* and sister species is covered by a sheet of extracellular matrix (Smith et al., 2020), which might be widespread in flies but species with asymmetric male genitalia may have a specific composition that is more permissive of mechanical force transmission among tissues. Since different parts of genitalia are asymmetric in closely related species of *D. pachea*, the diversification of asymmetry between the three *nannoptera* species may be associated with changes of extracellular matrix contacts, composition or deposition, that may target the lobes and phallus tip in *D. pachea*, just the phallus in *D. acanthoptera*, or the anal plates in *D. wassermani*. Under the signaling model (Figure 5B), the time period during which the growth signal is sent or received determines whether genitalia would form symmetric or asymmetric structures. If the signal is active throughout genitalia rotation, then both left and right sides receive the same amount of signal and develop similarly. If the signal is absent during part of the genitalia rotation, then the left and right sides will not receive as much signal and will acquire different sizes. This latter model implies that the timing and speed of male genitalia rotation relative to the timing of the growth signal is crucial for the determination of the final size of the lobes.

We took advantage of our *myo1D* mutant to investigate the relationship between genitalia asymmetry and right-sided mating posture. The proper dorso-ventral orientation of male genitalia was shown to be a prerequisite for copulation and efficient genitalia coupling in *D. melanogaster myo1D* mutants, with deviation from wild-type male genitalia orientation of less than 22° already tending to decrease reproduction and copulation success, both with wildtype and *myo1D* mutant females (Inatomi et al., 2019). We obtained similar results with *D. pachea myo1D*^*mut/mut*^ males that achieved copulation with a median deviation of only 15° from wild-type orientations and a maximum of 31° (Figure S6B). It corroborates the previous results that the proper alignment of female and male genital dorso-ventral axes are crucial in *Drosophila* to achieve a stable copulation position.

We found that *D. pachea* male genital asymmetry direction does not determine the sidedness of the mating posture since wildtype and mutant males with reversed genital asymmetry consistently mate in a right-sided copulation posture. Right-sided mating could therefore be determined by the female. We did not investigate mating posture in *myo1D*^*mut/mut*^ females. To our knowledge, no obvious left-right asymmetry has been described in female genitalia or genital tract in *D. pachea* (Rhebergen et al., 2016; van Gammeren et al., 2022). The complex of female and male genitalia during copulation of *D. pachea* has been investigated by microCT (van Gammeren et al., 2022). The asymmetric male phallus enters only the most distal part of the female uterus, which is composed of soft and deformable muscular tissue. The distal part of the female uterus surrounds and adopts the shape of the phallus and the anterior uterus lumen expands and re-arranges during copulation. Investigation of discrete uterus movement may help to test the possibility that the position of the male phallus, and with it the entire copulatory complex, might be actively controlled and moved by muscular movements of the female uterus. Changes of uterus shapes during copulation have also been reported for *D. melanogaster* (Mattei et al., 2015) and further investigations are needed to test their effect on mating position, sperm transfer and reproductive fitness in general. Alternatively, the right-sided mating posture of males might be a hard-wired behavior not depending on *Myo1D*. This would thus suggest the existence of other left-right determinants than *Myo1D* to determine lateralized behavior.

The right-sided mating posture could have possibly favored the evolution of morphological asymmetry as an instructive cue to optimize genital contacts and sperm transfer (Huber, 2010). However, mating behavior and evolution of genital morphology do not necessarily co-evolve. Lateralized copulation postures and asymmetric male genitalia have been described for a number of insects (Huber et al. 2007), such as in *Ciulfina* praying mantids (Holwell et al., 2015; Holwell and Herberstein, 2010), that reveal sinistral and dextral enantiomorphs of asymmetric male genitalia. However, both male enantiomorphs were found to adopt the same bending orientation during copulation of *Ciulfina baldersoni* (Heleodoro, 2022), indicating that this behavior does not influence the chirality of the genitalia. Mantids use spermatophore transfer instead of a penetrating phallus. This may enable a more flexible attachment of genitalia without a specific stereotyped positioning of the male phallus, as noted for *D. pachea* (Rhebergen et al., 2016; van Gammeren et al., 2022). It could possibly imply mating behavior or copulation positions to have a higher impact on the complex of female and male genitalia and efficient sperm transfer in species with internal fertilization because the mating posture directly affects the position of the phallus inside the female uterus. Evolution of genitalia morphology could therefore be a frequent consequence of changes in mating position in species with internal fertilization, although this hypothesis remains speculative.

Summarizing, the initial growth of *D. pachea* male genital lobes is synchronous with the clockwise 360° male genitalia rotation, which is a conserved tissue remodeling process in flies. The size ratio of the lobes depends on the completion of genitalia rotation in pyriproxyfen treated individuals and the sidedness of lobe asymmetry is associated with the direction of genitalia rotation in *D. pachea myo1D* mutants. Thus, genitalia rotation has likely been co-opted during evolution to provide developmental cues for lobe asymmetry direction and extent. Asymmetric lobe growth may depend on mechanical forces or signaling molecules that trigger different responses in the left and right sides due to clockwise tissue rotation. *D. pachea* mates in a right-sided copulation posture, but this is not a consequence of asymmetric male genitalia, since *myo1D* mutants with reversed genitalia asymmetry also mate in a right-sided position. Mating behavior is therefore determined by other cues and may depend on the female, or could be hard-wired in the male central nervous system. Altogether, our study reveals that a novel male genital lobe asymmetry has evolved through co-option of a pre-existing directional tissue remodeling process.

## Materials and methods

### Genitalia primordia preparation and immunohistochemistry

White pupae were transferred to moist tissue inside 5 cm petri dishes and left at 25°C inside plastic boxes for 24 h - 43 h. Pupae were then dissected by removing the pupal case, followed by transfer of the individual into PBS. The posterior part of the pupae was isolated and genitalia were gently detached from the cuticle by pumping PBS into the inside until genitalia floated out. Tissues were fixed at room temperature for 20 min in 4% PBS. Immunostaining was performed using a rabbit anti-PhosphoHistone H3 primary antibody (diluted at 1:200) (ThermoFisher #701258), a rabbit anti-cleaved Caspase 3 antibody (diluted at 1:400) (Cell Signaling Technology, #9661) and a secondary anti-rabbit-488 antibody (ThermoFisher, #SA5-10038) (diluted at 1:200). DAPI staining was then performed by a 15 min incubation in a 0.25 μg/mL solution (Sigma-Aldrich). Samples were mounted on a microscope slide with Vectashield® mounting medium. Confocal microscopy was performed on DMI8 (Leica) and ECLIPSE Ti2-E (Nikon) inverted microscopes, both equipped with a spinning-disc head CSU-W1 (Yokogawa Corporation of America) and controlled by MetaMorph software (Molecular Devices) and NIS-Elements software (Nikon), respectively. Nuclei and mitotic or apoptotic cells in lobes were quantified with Imaris software (Oxford Instruments, version 8.4.1) using spot and surface tools.

### Pyriproxyfen treatment

Pyriproxifen (Merck, #34174) stock solution of 0.1 mg/mL was diluted in acetonitrile according to Riddiford and Ashburner 1991, and serial dilutions were then prepared with acetone. *D. pachea* individuals undergoing puparium formation (white pupae) were aligned on a double-sided tesa stripe inside a 5 cm petri dish and 0.1 µL pyriproxyfen dilution was applied onto the posterior tip of each individual, corresponding to 0 pmol – 0.7 pmol. Petri-dishes with pupae were incubated in a moist plastic box at 25°C until the adults hatched. Adults were maintained at 25°C for one day and then sacrificed by transfer into absolute ethanol. Males were selected and imaged with a VHX2000 microscope (Keyence), followed by genitalia preparation, mounting in glycerol and imaging. Length and rotation measurements were performed with imageJ (Schindelin et al., 2012).

### Generation of *Myo1D* mutants

The *D. pachea myo1D* ortholog was identified by sequence comparisons of *D. melanogaster myo1D* cDNA sequence NM_001201855.2 and the *D. pachea* genome assembly JABVZX000000000 (Suvorov et al., 2021). CRISPR targets were searched manually and single guide RNA synthesis was performed *in vitro* according to (Bassett et al., 2013). The sgRNA/Cas-9 complexes were injected using standard Drosophila germline transformation procedures. The injection mix contained 0.1 μg/μL sgRNA, 2 μM NLS-Cas9 (New England Biolabs) and 1x NLS-Cas9 reaction Buffer (New England Biolabs). Emerging adults from injections were crossed to non-injected individuals and resulting progeny was maintained for several generations. Candidate stocks for CRISPR mutants were identified by visual inspection of male adults that had genitalia rotation defects. Candidate stocks were screened for *myo1D* mutations by PCR and Sanger sequencing. We identified a mutation at position 1724 downstream of the translation start codon, which contained a 12-bp deletion and a 7-bp insertion (Figure S2). The frameshift leads to a premature stop codon, abolishing 443 native codons that encode the C-terminal part of *Myo1D*. The identified mutation also contained a *Nco*I restriction site, not present in the wild-type allele. We took advantage of this restriction site to determine the genotype of *D. pachea* individuals. Prior to genotyping, genitalia of male adults or pupae were dissected and images were acquired. A detailed description of the procedures are given in the extended materials and methods (supplementary information).

### Generation act5C::DE-Cad-EYFP_2 flies

We generated a membrane specific EYFP cellular marker pB-act5C::DE-Cad-EYFP that contains a partial *D. melanogaster* DE-Cadherin (*shg*) and EYFP coding sequences (Figure S5). The assembled gene construct includes the *D. melanogaster shg* 5’UTR and two initial cadherin repeats, the DE-cadherin trans-membrane domain and a part of the cytoplasmic domain without the β-Catenin binding domain. The gene construct was inserted into the vector backbone of pBAC-ECFP-15xQUAS_TATA-mcd8-GFP-SV40 (Addgene 104878) and the 3xP3 ECFP integration reporter was replaced by 3xP3 DsRed (Matz et al., 1999) (Figure S5). transgenic flies were produced by germline transformation as described above. Identification of act5C::DE-Cad-EYFP_2 transgenic flies was achieved by evaluation of eye fluorescence driven by the 3xP3 DsRed integration reporter. The insertion site was mapped to an autosome at position 14,614,126 on contig tig00000094, inside the first 31.95-kb intron of the putative *D. pachea wheeler18* gene homolog. A detailed description of the procedures are given in the extended materials and methods (supplementary information).

### Genitalia live-imaging

We followed a particular crossing scheme (Figure S5) to analyze genitalia rotation of varying *myo1D* genotypes in a heterozygous *act5C::DE-Cad-EYFP_2* background. Time-lapse microscopy of developing genitalia was performed at 20 – 25 h APF on a DMI8 inverted microscope (Leica), equipped with a spinning-disc head CSU-W1 (Yokogawa Corporation of America) and controlled by MetaMorph software (Molecular Devices). Imaging up to 20 samples in parallel was carried out at 200-fold magnification for 15 -24 h, by acquisition of individual z-stacks every 30 min. Maximum z-projections were exported from each stack using imageJ and projections were concatenated to derive movies using standard command line functions in a bash terminal (Gnu, 2007).

### Analysis of mating behavior

We performed video-recording of virgin but sexually mature *D. pachea* individuals (Figure 4, Supplementary Dataset 6). Courtship and copulation duration analysis was carried out as described in (Lefèvre et al., 2021), and mating position analysis as in (Acurio et al., 2019). A detailed description of the procedures are given in the extended materials and methods (supplementary information).

### Statistics

Data analysis was done in R version 4.3.3. For Fig. 1, the 2- and 4-parameter logistic regressions and standard errors were calculated with the R-package ‘nplr’ (Commo and Bot, 2016). For Figure 2, logistic regression curves and confidence intervals were calculated with the R-package ‘drda’ (Malyutina et al., 2023).

## Acknowledgements

We thank Stéphane Prigent for preparation of Drosophila genitalia. We acknowledge the ImagoSeine core facility of the Institut Jacques Monod, member of IBiSA and France-BioImaging (ANR-10-INBS-04) infrastructures for assistance in microscopy. We thank Isabelle Germon, Véronique Borday-Birraux and Stecy Mienanzambi for help with immunohistochemistry experiments. We are grateful to Nicolas Gompel for initial help with *Drosophila* germline transfection and Allison J. Bardin and Carolina Parada for comments on previous drafts.

## Data availability

Data supporting this article has been deposited at the Zenodo database (DOI: 10.5281/zenodo.15394323, 10.5281/zenodo.15446323, 10.5281/zenodo.15446319, 10.5281/zenodo.15412039), except for DNA sequence data of the *D. pachea myo1D* coding sequence, which was deposited at NCBI (accession number OM240650).

## References

Acurio, A.E., Rhebergen, F.T., Paulus, S., Courtier-Orgogozo, V., Lang, M., 2019. Repeated evolution of asymmetric genitalia and right-sided mating behavior in the Drosophila nannoptera species group. BMC Evolutionary Biology 19, 109. 10.1186/s12862-019-1434-z

Ádám, G., Perrimon, N., Noselli, S., 2003. The retinoic-like juvenile hormone controls the looping of left-right asymmetric organs in Drosophila. Development 130, 2397–2406. 10.1242/dev.00460

Alsafwani, R.S., Nasser, K.K., Shinawi, T., Banaganapalli, B., ElSokary, H.A., Zaher, Z.F., Shaik, N.A., Abdelmohsen, G., Al-Aama, J.Y., Shapiro, A.J., 2021. Novel MYO1D Missense variant identified through whole exome sequencing and computational biology analysis expands the spectrum of causal genes of laterality defects. Frontiers in Medicine 8, 724826.

Bächli, G., Bernhard, U., Godknecht, A., 2021. TaxoDros [data base].

Bassett, A.R., Liu, J.-L., 2014. CRISPR/Cas9 and Genome Editing in Drosophila. Journal of Genetics and Genomics 41, 7–19. 10.1016/j.jgg.2013.12.004

Benitez, S.G., Sosa, C.M., Tomasini, N., Macías, A., 2010. Both JNK and apoptosis pathways regulate growth and terminalia rotation during Drosophila genital disc development. 10.1387/ijdb.082724sb

Blum, M., Ott, T., 2018. Animal left–right asymmetry. Current Biology 28, R301–R304.

Chougule, A., Lapraz, F., Földi, I., Cerezo, D., Mihály, J., Noselli, S., 2020. The Drosophila actin nucleator DAAM is essential for left-right asymmetry. PLOS Genetics 16, e1008758. 10.1371/journal.pgen.1008758

Commo, F., Bot, B.M., 2016. Package nplr: N-Parameter Logistic Regression. 2016. View Article.

Courtier-Orgogozo, V., Martin, A., 2020. The coding loci of evolution and domestication: current knowledge and implications for bio-inspired genome editing. Journal of Experimental Biology 223, jeb208934.

Coutelis, J.B., Petzoldt, A.G., Spéder, P., Suzanne, M., Noselli, S., 2008. Left–right asymmetry in Drosophila. Seminars in Cell & Developmental Biology, Cell Shape and Tissue Morphogenesis 19, 252–262. 10.1016/j.semcdb.2008.01.006

Eberhard, W.G., 2010. Evolution of genitalia: theories, evidence, and new directions. Genetica 138, 5–18.

Eberhard, W.G., 1985. Sexual selection and animal genitalia. Harvard University Press.

Feuerborn, H.J., 1922. Das Hypopygium “inversum” und “circumversum” der Dipteren. Zool. Anz 55, 89–213.

Géminard, C., González-Morales, N., Coutelis, J.-B., Noselli, S., 2014. The myosin ID pathway and left–right asymmetry in Drosophila. genesis 52, 471–480. 10.1002/dvg.22763

Gnu, P., 2007. Free software foundation. Bash (3. 2. 48)[Unix shell program] 624.

González-Morales, N., Géminard, C., Lebreton, G., Cerezo, D., Coutelis, J.-B., Noselli, S., 2015. The Atypical Cadherin Dachsous Controls Left-Right Asymmetry in Drosophila. Developmental Cell 33, 675–689. 10.1016/j.devcel.2015.04.026

Hamada, H., Tam, P., 2020. Diversity of left-right symmetry breaking strategy in animals. F1000Res 9, F1000 Faculty Rev-123. 10.12688/f1000research.21670.1

Heleodoro, R.A., 2022. The first two cases of antisymmetry in the male genitalia of Phasmatodea reveal a new species of Isagoras Stål, 1875 (Phasmatodea: Pseudophasmatidae: Xerosomatinae) from the Brazilian Atlantic Forest. Zoologischer Anzeiger 296, 161–178.

Holwell, G.I., Herberstein, M.E., 2010. Chirally dimorphic male genitalia in praying mantids (Ciulfina : Liturgusidae). Journal of Morphology 271, 1176–1184. 10.1002/jmor.10861

Holwell, G.I., Kazakova, O., Evans, F., O’Hanlon, J.C., Barry, K.L., 2015. The functional significance of chiral genitalia: Patterns of asymmetry, functional morphology and mating success in the praying mantis Ciulfina baldersoni. PLoS One 10, e0128755.

Horn, C., Offen, N., Nystedt, S., Häcker, U., Wimmer, E.A., 2003. piggyBac-based insertional mutagenesis and enhancer detection as a tool for functional insect genomics. Genetics 163, 647–661.

Hozumi, S., Maeda, R., Taniguchi, K., Kanai, M., Shirakabe, S., Sasamura, T., Spéder, P., Noselli, S., Aigaki, T., Murakami, R., Matsuno, K., 2006. An unconventional myosin in Drosophila reverses the default handedness in visceral organs. Nature 440, 798–802. 10.1038/nature04625

Huber, B.A., 2010. Mating positions and the evolution of asymmetric insect genitalia. Genetica 138, 19–25.

Huber, B.A., Sinclair, B.J., Schmitt, M., 2007. The evolution of asymmetric genitalia in spiders and insects. Biological Reviews 82, 647–698. 10.1111/j.1469-185X.2007.00029.x

Inatomi, M., Shin, D., Lai, Y.-T., Matsuno, K., 2019. Proper direction of male genitalia is prerequisite for copulation in Drosophila, implying cooperative evolution between genitalia rotation and mating behavior. Sci Rep 9, 210. 10.1038/s41598-018-36301-7

Jinek, M., Chylinski, K., Fonfara, I., Hauer, M., Doudna, J.A., Charpentier, E., 2012. A Programmable Dual-RNA–Guided DNA Endonuclease in Adaptive Bacterial Immunity. Science 337, 816–821. 10.1126/science.1225829

Juan, T., Géminard, C., Coutelis, J.-B., Cerezo, D., Polès, S., Noselli, S., Fürthauer, M., 2018. Myosin1D is an evolutionarily conserved regulator of animal left–right asymmetry. Nat Commun 9, 1942. 10.1038/s41467-018-04284-8

Lai, Y.-T., Sasamura, T., Kuroda, J., Maeda, R., Nakamura, M., Hatori, R., Ishibashi, T., Taniguchi, K., Ooike, M., Taguchi, T., 2023. The Drosophila AWP1 ortholog Doctor No regulates JAK/STAT signaling for left–right asymmetry in the gut by promoting receptor endocytosis. Development 150, dev201224.

Lang, M., Orgogozo, V., 2012. Distinct copulation positions in Drosophila pachea males with symmetric or asymmetric external genitalia. Contributions to Zoology 81.

Lang, M., Polihronakis Richmond, M., Acurio, A.E., Markow, T.A., Orgogozo, V., 2014. Radiation of the Drosophila nannoptera species group in Mexico. Journal of evolutionary biology 27, 575–584.

Lebreton, G., Géminard, C., Lapraz, F., Pyrpassopoulos, S., Cerezo, D., Spéder, P., Ostap, E.M., Noselli, S., 2018. Molecular to organismal chirality is induced by the conserved myosin 1D. Science 362, 949–952.

Lefèvre, B.M., Catté, D., Courtier-Orgogozo, V., Lang, M., 2021. Male genital lobe morphology affects the chance to copulate in Drosophila pachea. BMC ecology and evolution 21, 1–13.

Macias, A.M., Romero, N.M., Martín, F., Suárez, L., Rosa, A.L., Morata, G., 2004. PVF1/PVR signaling and apoptosis promotes the rotation and dorsal closure of the Drosophila male terminalia. 10.1387/ijdb.041859am

Malyutina, A., Tang, J., Pessia, A., 2023. drda: An R package for dose-response data analysis using logistic functions. Journal of Statistical Software 106, 1–26.

Mattei, A.L., Riccio, M.L., Avila, F.W., Wolfner, M.F., 2015. Integrated 3D view of postmating responses by the Drosophila melanogaster female reproductive tract, obtained by micro-computed tomography scanning. Proc. Natl. Acad. Sci. U.S.A. 112, 8475–8480. 10.1073/pnas.1505797112

Palmer, A.R., 2009. Animal asymmetry. Current Biology 19, R473–R477. 10.1016/j.cub.2009.04.006

Petzoldt, A.G., Coutelis, J.-B., Géminard, C., Spéder, P., Suzanne, M., Cerezo, D., Noselli, S., 2012. DE-Cadherin regulates unconventional Myosin ID and Myosin IC in Drosophila left-right asymmetry establishment. Development 139, 1874–1884.

Pitnick, S., Heed, W.B., 1994. New species of cactus-breeding Drosophila (Diptera: Drosophilidae) in the nannoptera species group. Annals of the Entomological Society of America 87, 307–310.

Rhebergen, F.T., Courtier-Orgogozo, V., Dumont, J., Schilthuizen, M., Lang, M., 2016. Drosophila pachea asymmetric lobes are part of a grasping device and stabilize one-sided mating. BMC evolutionary biology 16, 176.

Rice, G., David, J.R., Kamimura, Y., Masly, J.P., Mcgregor, A.P., Nagy, O., Noselli, S., Nunes, M.D.S., O’Grady, P., Sánchez-Herrero, E., Siegal, M.L., Toda, M.J., Rebeiz, M., Courtier-Orgogozo, V., Yassin, A., 2019. A standardized nomenclature and atlas of the male terminalia of Drosophila melanogaster. Fly 13, 51–64. 10.1080/19336934.2019.1653733

Riddiford, L.M., Ashburner, M., 1991. Effects of juvenile hormone mimics on larval development and metamorphosis of Drosophila melanogaster. General and comparative endocrinology 82, 172–183.

Rousset, R., Bono-Lauriol, S., Gettings, M., Suzanne, M., Spéder, P., Noselli, S., 2010. The Drosophila serine protease homologue Scarface regulates JNK signalling in a negative-feedback loop during epithelial morphogenesis. Development 137, 2177–2186. 10.1242/dev.050781

Saydmohammed, M., Yagi, H., Calderon, M., Clark, M.J., Feinstein, T., Sun, M., Stolz, D.B., Watkins, S.C., Amack, J.D., Lo, C.W., 2018. Vertebrate myosin 1d regulates left–right organizer morphogenesis and laterality. Nature Communications 9, 3381.

Schilthuizen, M., 2014. Nature’s nether regions. Viking, New York.

Schilthuizen, M., 2013. Something gone awry: unsolved mysteries in the evolution of asymmetric animal genitalia. Animal Biol 63, 1–20. 10.1163/15707563-00002398

Schindelin, J., Arganda-Carreras, I., Frise, E., Kaynig, V., Longair, M., Pietzsch, T., Preibisch, S., Rueden, C., Saalfeld, S., Schmid, B., 2012. Fiji: an open-source platform for biological-image analysis. Nature methods 9, 676–682.

Smith, S.J., Davidson, L.A., Rebeiz, M., 2020. Evolutionary expansion of apical extracellular matrix is required for the elongation of cells in a novel structure. eLife 9, e55965. 10.7554/eLife.55965

Spéder, P., Ádám, G., Noselli, S., 2006. Type ID unconventional myosin controls left–right asymmetry in Drosophila. Nature 440, 803–807.

Stern, D.L., 2022. Transgenic tools for targeted chromosome rearrangements allow construction of balancer chromosomes in non-melanogaster Drosophila species. G3 12, jkac030.

Suvorov, A., Kim, B.Y., Wang, J., Armstrong, E.E., Peede, D., D’agostino, E.R., Price, D.K., Waddell, P.J., Lang, M., Courtier-Orgogozo, V., 2022. Widespread introgression across a phylogeny of 155 Drosophila genomes. Current Biology 32, 111–123.

Suzanne, M., Petzoldt, A.G., Spéder, P., Coutelis, J.-B., Steller, H., Noselli, S., 2010. Coupling of apoptosis and L/R patterning controls stepwise organ looping. Current Biology 20, 1773–1778.

Taniguchi, K., Hozumi, S., Maeda, R., Ooike, M., Sasamura, T., Aigaki, T., Matsuno, K., 2007. D-JNK signaling in visceral muscle cells controls the laterality of the Drosophila gut. Developmental Biology 311, 251–263. 10.1016/j.ydbio.2007.08.048

Taniguchi, K., Maeda, R., Ando, T., Okumura, T., Nakazawa, N., Hatori, R., Nakamura, M., Hozumi, S., Fujiwara, H., Matsuno, K., 2011. Chirality in Planar Cell Shape Contributes to Left-Right Asymmetric Epithelial Morphogenesis. Science 333, 339–341. 10.1126/science.1200940

Tingler, M., Kurz, S., Maerker, M., Ott, T., Fuhl, F., Schweickert, A., LeBlanc-Straceski, J.M., Noselli, S., Blum, M., 2018. A conserved role of the unconventional myosin 1d in laterality determination. Current Biology 28, 810–816.

van Gammeren, S., Lang, M., Rücklin, M., Schilthuizen, M., 2022. No evidence for asymmetric sperm deposition in a species with asymmetric male genitalia. PeerJ 10, e14225.

Vilela, C.R., Bächli, G., 1990. Taxonomic studies on Neotropical species of seven genera of Drosophilidae (Diptera).

