## Supplementary Materials for "Evolution of a novel left-right asymmetry in organ size by co-option of a tissue rotation process"

[2]current address: Team "Stem Cells and Tissue Homeostasis", Institut Curie, CNRS, UMR3215, INSERM U934, PSL Research University, 26 rue d'Ulm, 75248 Paris Cedex 05

[3]current address: Laboratoire Évolution, Génomes, Comportement et Écologie, CNRS, IRD, Université Paris-Saclay – Institut Diversité, Ecologie et Evolution du Vivant (IDEEV), 91190 Gif-sur-Yvette, France

**This PDF file includes:**

Extended Materials and Methods

Supplementary Figures S1 to S10

Supplementary Table S1

Legends for Movies 1 to 2 and Datasets S1 to S8

SI References

### Extended Materials and Methods

#### *Drosophila* maintenance.

We maintained *D. melanogaster* stock T.7 (Tucson Drosophila Species Stock Center, ref. 14021-0231.7) and *D. pachea* inbred line 14.2 (11 generations of single couple mating of stock 15090-1698.01 from Drosophila Species Stock Center) in 25 x 95 mm plastic vials containing 10 mL of standard Drosophila medium (60 g/L brewer's yeast, 66.6 g/L cornmeal, 8.6 g/L agar, 5 g/L methyl-4-hydroxybenzoate and 2.5% v/v ethanol) and a ~ 10 x 50 mm piece of bench protection sheet (Bench guard). *D. pachea* cannot convert dietary cholesterol into 7-dehydrocholesterol for ecdysone hormone synthesis (Heed and Kircher, 1965). We mixed the medium of each vial with 40 µL of 5 mg/mL 7-dehydrocholesterol (Sigma-Aldrich 30800, Merck dissolved in ethanol) to enable cultivation of this species in the laboratory. Flies were kept at 25°C inside incubators (Velp) at constant light (approximately 200 lumen). For behavioral assays, flies were kept at a 12h light: 12h dark photo-periodic cycle combined with a 30-min linear illumination change between light (1080 lumen) and dark.

#### Preparation of CRISPR single guide RNA.

The *D. pachea* ortholog of *D. melanogaster myo31df* gene (here named *myo1D*) was identified by local BLAST using *D. melanogaster* m*yo1D* coding sequence (GenBank Accession number NM_001201855.2) as a query on contig tig00000030 of the newly derived *D. pachea* genome assembly (Genbank accession JABVZX000000000) (Suvorov et al., 2022). The region of the predicted coding sequence was first deduced from sequence alignments with the *D. melanogaster* m*yo1D* coding sequence. Then, *D. pachea myo1D* coding sequence of stock 15090-1698.01 was determined (GenBank accession number OM240650) by Sanger sequencing of cDNA from total RNA, extracted from 5 females and 5 males with the Nucleospin RNA mini kit (Macherey-Nagel). First strand synthesis was carried out with the SuperScript III First-Strand Synthesis kit (ThermoFisher), using 1 µg total RNA and 1.2 µM of oligonucleotide pacMyo1D-retro (Table S1) in 25 µL reaction volume and incubation for 2 hours at 55 °C with an initial denaturation step for 5 min at 65 °C before addition of SuperScript III. The *myo1D* coding sequence was then amplified by PCR with oligonucleotides pacMyo1D-debut and pacMyo1D-stop (Table S1) that covered the putative translation start and the predicted stop codon. PCR product was sent to Eurofins for Sanger sequencing with oligonucleotides pacMyo1D-debut, pacMyo1D-R4, Dpa2Rbis3, 2F_Dpac_m31DF, and 4F_Dpac_m31DF (Table S1). All PCR reactions were carried out in the thermocycler iCycler (BioRad) and sequence data was analyzed with the program Geneious 6.0.6 (www.geneious.com). The putative *D. pachea myo1D* coding sequence revealed a length of 3036 bp and contained nine exons (Figure S3 A). The design and preparation of single guide RNAs (sgRNAs) (Table S1) was performed according to (Bassett et al., 2013). Briefly, DNA target $GGN_{19}GG$ was identified at nucleotide positions 1710 - 1732 in the coding region of the putative *D. pachea myo1D* homolog, corresponding to positions 2,340, 437 - 2,340,459 of contig tig00000030 of the *D. pachea* genome assembly (Figure S3 A). The first 20 nucleotides of this target were identical to the gene specific sgRNA sequence motif and the last three nucleotides formed the protospacer adjacent motif (PAM) in the *myo1D* gene (Figure S3 B). The DNA target was assembled to the *D. pachea* genome draft using bowtie 1.1.22 (Langmead et al., 2009) to search for potential off-target sequences and only revealed alignments with at least three nucleotide substitutions. Production of sgRNA was prepared with oligonucleotides Dpa_myo1DCRISPR_F and sgR (Table S1). PCR products were spin-column purified with the PCR clean-up kit (Macherey & Nagel) and 1.5 - 2.0 µg were used as template for *in vitro* transcription and purification of sgRNA with the T7 MEGAscript kit (Ambion). Final sgRNA were stored at -80°C, at a concentration of 1 µg/µL. The CRISPR/Cas-9 injection mix for germline transformation contained 0.1 µg/µL sgRNA, 2 µM NLS-Cas9 (New England Biolabs) and 1x NLS-Cas9 reaction Buffer (New England Biolabs).

**Generation of pB-act5C::DE-Cad-EYFP.**

We generated a membrane specific EYFP cellular marker construct that contains a partial *D. melanogaster* DE-Cadherin (*shg*) coding sequence (CDS) of stock T.7 (Figure S5 A). We PCR-amplified two fragments of the *shg* CDS, corresponding to nucleotide positions -724 - 903 (Fragment_1) and 3985 - 4326 (Fragment_2) relative to the annotated start codon in sequence NM_057374 (Genbank Accession Number). Our aim was to include coding sequence parts that encode domains important for protein localization to the cell membrane while annotated protein-interaction domains were avoided. Fragment_1 encodes the *D. melanogaster shg* 5'UTR and two initial cadherin repeats that are majorly cleaved as a DE-cadherin specific pro-peptide (Oda et al., 1994; Pacquelet et al., 2003). Fragment_2 corresponds to the DE-cadherin trans-membrane domain and a part of the cytoplasmic domain, but lacks the β-Catenin binding domain. We extracted total RNA from 10 adult flies with the Nucleospin RNA mini kit (Macherey-Nagel). First strand synthesis of *shg* cDNA was carried out as described above with the SuperScript III First-Strand Synthesis kit (ThermoFisher), using 5 µg total RNA and 1.2 µM of oligonucleotide m_DECad_R1 (Table S1). Fragment_1 and Fragment_2 were amplified with the Phusion High-Fidelity PCR Kit (New England Biolabls) from the cDNA with oligonucleotides m_DECad_F1 / mDECad_R2 and m_DECad_F2 / m_DECad_R3, respectively (Table S1). Oligonucleotides were designed to allow Gibson cloning (Gibson et al., 2009) and contained an overlap of 12-20 nucleotides with respect to adjacent DNA fragments and the pGEM®-T Easy cloning vector (Promega). Gibson Assembly was performed according to (Gibson et al., 2009), except that the assembly reactions were incubated for 10 min at 37°C and then for 3 hr at 50°C, as described in (Nagy et al., 2018). We used 1 µL of assembly mixture for chemical transformation of constructs into 25 µL NEB 10-beta (New England Biolabs) competent cells. Ampicillin-resistant colonies were selected on 100 mg/mL Amp-LB plates. The components for the Gibson Assembly Master-mix (Gibson et al., 2009) were purchased from Sigma-Aldrich (now Merck). We used standard thermocycler programs, recommended by the manual of the kit and oligonucleotide specific annealing temperatures (Table S1). PCR products were spin-column purified with the PCR clean-up kit (Macherey & Nagel). Fragment_1 was then inserted upstream of Fragment_2 into the pGEMT cloning vector and a single joint fragment was cloned by Gibson cloning into *Xho*I linearized plasmid act::EYFP, which is a modified version of 3XP3::EYFP vector (Horn et al., 2003) (Figure S5 B). It contains a single *Xho*I restriction site between the *D. melanogaster* actin 5C promoter (Thummel et al., 1988) (upstream) and the EYFP coding sequence with a SV40 terminator (Horn et al., 2003). The resulting clone act5C::DE-Cad-EYFP_1 revealed variable lengths in some preparations due to partial loss of the EYFP coding sequence. Therefore, we amplified the region act::DE-cadherin-EYFP-SV40 in two fragments (Fragment_3 and Fragment_4) by PCR with the Phusion High-Fidelity PCR Kit (NEB) using oligonucleotides act5C_F1 and act5C_R and act5C_F2 and SV40_R (Table S1) and inserted them by Gibson cloning into the vector backbone of pBAC-ECFP-15xQUAS_TATA-mcd8-GFP-SV40 (Addgene 104878), digested with restriction enzymes *Nde*I and *Bgl*II. In addition, we exchanged the 3xP3 ECFP integration reporter gene by 3xP3 DsRed (Matz et al., 1999). The integrity of the final construct act5C::DE-Cad-EYFP_2 (Figure S5 A) was verified by Sanger sequencing. The final injection mix for germline transformation contained 150 ng/µL piggyBac helper plasmid (Horn et al., 2003), 150 ng/µL act5C::DE-Cad-EYFP_2 and 1x NLS-Cas9 reaction Buffer (New England Biolabs).

**Germline transformation (for production of transgenic strains and CRISPR mutants).**

About 250-500 adult *D. pachea* flies were maintained at 25°C and constant light inside custom-made cylindrical egg-laying cages (6 cm x 8 cm, diameter x height), closed at the bottom with a 6-cm petri dish containing grape juice agar: 24 gr / L agar, 26.4 gr / L sucrose, 1/5 volume grape juice, 1.2 gr / L Tegosept and 0.02 gr / L 7-dehydrocholesterol (7DHC), with about 200 µL yeast paste (fresh baker yeast) on the surface. Eggs were collected in 2-hour intervals by exchanging the feeding plates, by washing the plate surface with temperate tap water, which was then passed through a 100 micron nylon filter (BD Falcon 352360). Eggs were dechorionated by strong agitation of the filter for 90 sec in 1.3% bleach (BEC) and then rinsed extensively with running water. Eggs were then aligned on a 5 mm thick piece of 2% agar (prepared with tap water) under a stereomicroscope K-500 (VWR) with the help of fine forceps and attached to a 18x18 mm microscopy cover slip (Menzel-Gläser, VWR) coated with TESA glue. For glue coating, about 50 cm of double-sided transparent tesafilm® (Tesa) was dissolved in 20 mL n-heptane and 15 µL of this solution was pipetted onto a ~ 3 mm wide stripe at one edge of the cover slip. After evaporation of n-heptane, embryos were lifted with the glue-coated side of the cover slip from the agar and allowed to dry for 5 min at room temperature before being covered with 40 µL (2 drops) halocarbon oil Voltalef 10S (VWR). Early embryos at the stage of the syncytial blastoderm were injected with injection mix into the posterior pole. The amount of mix corresponded to a drop of approximately 1/3 the volume of the convex posterior end of the embryo. The manipulation was carried out on a light microscope Leica DM LS (Leica) at 100 fold magnification, with 1-mm borosilicate capillaries GB 100-50-10 (WPI) prepared with a needle puller P1000 (Sutter, parameters: heat 458, pull 70, speed 80, delay 200, pressure 500) that were connected to a micromanipulator 056530 (Leitz) and a micro-injector Transjector 5246 (Eppendorf). Upon injection, embryos were pushed towards the needle until it entered the posterior pole of the embryo. After injection, embryos were let to develop at 25°C for about 32 h. Then, larvae were collected from the halocarbon oil with fine forceps and were transferred to a 3.5 cm petri dish containing 1.2 gr / L potato starch (Mousseline, Nestlé), mixed with 1.2 gr / L Tegosept (USBiological) and 0.02 gr / L 7-dehydrocholesterol (Sigma-Aldrich 30800, Merck). Petri dishes were covered with a 3D-printed cylindrical lid of 3.5 cm x 2 cm (diameter x height) covered with a 100 micron nylon mesh and flies were let to develop inside, at 25°C and saturated humidity.

**Identification of *myo1D* mutants.**

Virgin adults from the injection trials were collected and isolated at 0-24 h after emerging from the pupa and crossed with 3 individuals of the opposite sex of the inbred line (from stock 15090-1698.01). Single progeny individuals were then either crossed back to 3 individuals of the opposite sex of the inbred line (from stock 15090-1698.01) or were pooled into single vials (2-5 females plus 2-5 males). These crosses were then maintained as stocks. To identify CRISPR mutants, adult males of consecutive generations were screened under a binocular microscope for genitalia rotation defects, characteristic for *myo1D* mutants in *D. melanogaster* (Spéder et al., 2006). Genomic DNA was isolated from single individuals with the DNeasy Blood & Tissue Kit (QIAGEN) or the Insect DNA kit (EZNA). The genomic region containing the CRISPR target site was amplified by PCR as described above with the Phusion High-Fidelity PCR Kit (New England Biolabs) in 35 µL reaction volume, using ~ 10 ngr genomic DNA and oligonucelotides 1F_Dpac_m31DF / Dpa2Rbis3, (Table S1). PCR products were sent to Eurofins for Sanger sequencing. We identified CRISPR mutants in the progeny of a single injected individual out of 1381 injected embryos in total. Mutants contained a 13-bp deletion around the PAM site and a 7-bp insertion. The mutation causes a frameshift and a premature stop codon 73 codons downstream of the induced mutation. It abolishes 443 native codons that correspond to the C-terminal part of the myosin motor domain, the calmodulin binding domain, and membrane diffusion domain (procite scan, https://prosite.expasy.org/prosite.html). The identified mutation also contained a *Nco*I restriction site, not present in the wild-type allele. We used this restriction site to determine the genotype of *D. pachea* individuals. For this, CRISPR target sites were amplified with the Phusion High-Fidelity PCR Kit (New England Biolabs) and oligonucleotides Myo1D_3F-genoF1/Dpa2Rbis3 in 20µL reaction volume. For amplifications, we used a touch-down PCR thermocycler program with the annealing temperature of the first ten cycles decreasing from 70 °C to 60°, followed by 25 cycles with a constant annealing temperature of 60 °C (Table S1). A total of 10 µL PCR product was mixed with 10 µL of *Nco*I dilution, containing 1x fastdigest Buffer (Thermo) and 0.5 µL *Nco*I-HF (New England Biolabs). Restriction digestion was carried out for 1h at 37 °C and finally DNA fragment lengths were separated by gel electrophoresis on a 2% agarose gel containing SYBR™ Safe DNA Gel Stain (ThermoFisher) in 1x Tris-Acetate-EDTA (TAE) Buffer and run at 3-4 Volt/cm next to a well containing 500 ng of 100 bp-ladder (New England Bioloabs). Gels were examined on a

UV-transilluminator Ebox VX2 (Vilber-Lourmat). Undigested DNA fragments of 215 bp corresponded to the wild-type *myo1D* allele whereas the mutant allele was cut into two fragments of 166 bp and 40 bp, neglecting NcoI 5' overhangs.

**Dissection and imaging of male specimens.**

Genitalia of male adults or pupae that had developed sclerotized genital tissue were dissected and imaged using a VHX 2000 microscope (Keyence), equipped with a 100-1000x VH-Z100W (Keyence) zoom objective. Individuals were fixed onto a dissection dish with fine needles with the ventral abdomen facing to the camera objective. An image of the entire body was acquired at 100 fold magnification to monitor the orientation angle of male genitalia relative to the male antero-posterior midline (Figure S2 B) of the abdomen was isolated. Pupae were treated as adults (Figure S2 C). The rest of the body was stored in 96% ethanol for DNA extraction, while genitalia were rehydrated in water for several minutes. Genitalia were dissected out with fine forceps and transferred into a transparent dish filled with glycerol for imaging at 400 fold magnification. Samples were leveled with respect to the microscope objective so that the basis of each left and right lobe, and the dorsal edge of the genital arch would be in the same focal plane. Dissected tissues were stored at 4 °C in Glycerol:Acetate:Ethanol (1:1:2). Orientation and length measurements were taken on acquired images using ImageJ version 1.50d (https://imagej.nih.gov/ij). A general trend towards a shortening of the abdomen is observed in Brachyceran Diptera, especially in Muscomorpha, and more pronounced in males than in females (McAlpine, 1981). In *Drosophila* males, this results in a location of the external genitalia at the ventral body side and the dorso-ventral axis of male genitalia approximates the overall antero-posterior axis of the abdomen. Genitalia orientation was estimated as the angle between these two axes when the animal is viewed from the ventral side (Figure S3 B,C), approximated by: 1) a line from the dorsal midpoint of the male anal plates, passing through the midpoint between male claspers, 2) a line from the dorsal midpoint of the anal plates towards the thorax, but parallel to the male midline, which was estimated by the medially located male sternites and the basis of the legs (Figure S3 B). In pupa, the midline axis was approximated by the location of the ventrally located legs (Figure S3 C). In previous studies (*3-5*), lobe lengths were measured as the distance between the base of a lateral spine located at the base of each lobe, and the medial tip of each lobe (Figure 2 F). However, we could not identify the lateral spines in some individuals with partially rotated genitalia, indicating that rotation progress is essential for the development of these spines. Alternatively, lateral spines might have been lost during dissection. The paired surstili of *D. pachea* have a rhomboid shape with an apical bud at their outer corner (Figure 2 F). In this study, we used this bud of each left and right surstylus as a landmark to measure the distal lobe length from the bud to the tip of each lobe. For phallus dissections, genitalia were boiled for 10 min in 30% KOH and then the left and right junctions of the hypandrium (internal genitalia) were cut. The phallus was oriented inside a transparent plastic dish filled with glycerol with the phallus tip pointing upwards and the hypandrium junction towards the Keyence VHX 2000 microscope objective (see above). This resulted in a ventral view of the phallus (Figure 3 F). Images were acquired at 600 fold magnification.

**Identification of act5C::DE-Cad-EYFP_2 transgenic flies.**

We injected the piggyBac construct act5C::DE-Cad-EYFP_2 into embryos of the *D. pachea myo1D* mutant. Emerging adults were crossed to 3 individuals of opposite sex of the *myo1D* mutant stock and transgenic progeny individuals were identified based on 3xP3::DsRed fluorescence in the eye (Horn et al., 2003), using a fluorescence stereomicroscope (Nikon SMZ 1500) equipped with a pE-300 (coolLED) illumination system and DsRed ET Filter set (AHF Analysetechnik). We identified transgenic individuals only from the progeny of a single injected individual out of 450 injected embryos. Transgenic flies were crossed to males of the *myo1D* mutant stock and transgenic progeny was selected by red-eye fluorescence in consecutive crosses to establish the *myo1D* mutant stock, used for further experiments. We identified the insertion site of the piggybac vector by inverse PCR (Ochman et al., 1988). For this, 10 adult flies were pooled and genomic DNA was extracted as described above. Genomic DNA was digested with the restriction enzymes SpeI and *Xba*I, or only SpeI, and subsequently purified with the PCR cleanup kit (Macherey and Nagel). We self-ligated purified DNA at dilutions of 0.25 ng/µL and 0.5 ng/µL overnight at 16°C in 10 µL reaction volume containing 1x ligation buffer and 1 µL T4 DNA ligase (ThermoFisher). A total of 1 µL of ligation product was used as DNA template in PCR amplifications with the Phusion taq kit (NEB), following the manufacturer's instructions for the reaction components and using oligonucleotides InvPCR_F1/InvPCRF2, InvPCR_R1/InvPCRR2 (supplementary dataset S9) in 25 µL reaction volume. Oligonucleotides were specific to the left arm piggyBac inverted repeat region. Purified PCR products were sent for Sanger sequencing and sequence data was aligned to the *D. pachea* genome assembly. The insertion site was mapped to position 14,614,126 on contig tig00000094. This position is inside the first 31.95-kb intron of the putative *D. pachea wheeler18* gene homolog, 26.417 kb downstream and 5.533 kb upstream of the 5'- and 3' splice sites (Figure S5 C). For time-lapse imaging experiments, we followed a particular crossing scheme in order to analyze genitalia rotation of varying *myo1D* genotypes in a heterozygous act5C::DE-Cad-EYFP_2 background (Figure S6). Females [$myo1D^{mut/mut}$, act5C::DE-Cad-EYFP_$2^{+/+}$] were crossed to wildtype males [$myo1D^{wt/wt}$, act5C::DE-Cad-EYFP_$2^{-/-}$]. The female progeny was then crossed back to wildtype males and male progeny was crossed to *myo1D* mutant males [$myo1D^{mut/mut}$, act5C::DE-Cad-EYFP_$2^{-/-}$]. Progeny females [act5C::DE-Cad-EYFP_$2^{+/-}$] of the latter cross were then crossed with non-fluorescent sibling males [act5C::DE-Cad-EYFP_$2^{-/-}$] and vice versa non-fluorescent females [act5C::DE-Cad-EYFP_$2^{-/-}$] were crossed with fluorescent males [act5C::DE-Cad-EYFP_$2^{+/-}$]. The presence of the act5C::DE-Cad-EYFP_2 insertion could be followed by the 3xP3::DsRed integration marker of the construct, visible as red fluorescence in the adult eyes or a red fluorescent central nervous system in larvae and early developing pupae.

**Time lapse microscopy.**

Live-imaging of developing genitalia was performed with about 20 pupae per microscopy session and was initiated at 20h-25h after the puparium had formed. The posterior part of each pupal case was removed with forceps (Dumont) and pupae were subsequently placed with the posterior end down into 1-mm holes inside a layer of solid, 2.5 mL 1% agarose, which was casted onto a 32 mm (diameter) round cover slip (0.17 mm, ThermoScientific) inside a circular POC-R2 Cell Cultivation System (PECON). The posterior end of each pupa was verified by eye to touch the cover slip. Moist tissue paper was put on top of the agarose along the PeCon cell wall and the cell was covered with a lid of a 3.5 cm plastic petri dish to prevent as much as possible dehydration of the agarose. We placed the pupae in an asymmetric arrangement into the cell and labeled each side of the cell in order to avoid sample confusion. Pupae were then examined on a DMI8 inverted microscope (Leica), equipped with a spinning-disc head CSU-W1 (Yokogawa Corporation of America) and controlled by MetaMorph software (Molecular Devices). Time lapse imaging was carried out at 200 fold magnification for 15-24 hours, by acquisition of individual z-stacks of 40-60 frames and a step size of 4 µm, using a 488 nm laser at 75% intensity, a 500-550 nm emission filter and 650 ms exposure time. Maximum z-projections were exported from each stack using a imageJ macro and projections were automatically concatenated to derive movies with conventional bash scripts. After microscopy, pupae were transferred into fresh 1% agarose and kept inside a plastic box with moist tissue paper on the bottom. Pupae were observed once every 24 hours. Specimens were considered to have died and were transferred into 96% ethanol if no developmental progress of metamorphosis was visible according to morphological markers as established by (Bainbridge and Bownes, 1981). Otherwise, individuals were let to develop to adulthood and then transferred into 96% ethanol.

**Copulation recording.**

Freshly emerged flies (0-3 days) were anesthetized with $CO_2$, separated according to sex and transferred into food vials in groups of 5-15 females or males using a Stemi 2000 (Zeiss) stereomicroscope, a $CO_2$-pad (Inject+Matic sleeper) and a brush. Flies were maintained at 25°C until they reached sexual maturity, at least 4 days for females and at least 14 days for males (Pitnick, 1993). Males were isolated into single vials for at least 2 days before the experiment was performed. Video recording was performed as described in (Acurio et al., 2019) and (Lefèvre et al., 2021). Briefly, one male and one female were introduced into a circular plastic mating cell with a diameter of 20 mm, a depth of 4 mm and a transparent 1-mm Plexiglas cover. Movies were recorded in a climate controlled chamber at 25°C ± 0.1°C and 80% ± 5% humidity. Flies were filmed from above using a monochrome camera Chameleon 3 (FLIR), equipped with a 50 mm objective (Thorlabs), using a cold light source for illumination from the side. Movies were recorded with FlyCapture2 SDK (FLIR) at a resolution of 1200 X 840 pixels. Movies were recorded until copulation ended or for at least 60 min when no copulation was detectable. Flies were then stored in 96% ethanol at -20°C.

**Mating position analysis.**

Movies were analyzed with the video editor OpenShot 1.4.3 (Open Shot Studios, Texas, USA). Courtship start, copulation start, the settling time point and the end of copulation were annotated manually as described in (Acurio et al., 2019) and (Lefèvre et al., 2021). Courtship was defined to start when the male displayed at least three consecutive typical male courtship behaviors (Spieth, 1952), such as tapping the female abdomen with the legs, wing vibration, following the female, and male licking behavior where the male touches the female ovipositor or the ground next to it with the proboscis (mouth-parts). This latter behavior was easy to spot and was monitored until copulation occurred (Figure S7). Courtship was defined to end with the start of copulation, when a male had mounted the female abdomen and had settled into an invariant copulation posture. Upon mounting, the male moves the abdomen and legs and the time from mounting until being settled for at least 30 seconds was annotated as mounting attempt (Figure S7). Males that did not settle kept moving on the female's abdomen or descended from the female. Copulation was defined to end when the male had completely descended from the female abdomen with the forelegs detached from the female dorsum, and female and male genitalia no longer in contact. We video-recorded 135 mating trials, of which 63 were used for comparison of courtship and copulation (supplementary dataset S6, Figure S7). We excluded 54 trials because no courtship or copulation was observed within 1 h of recording, 3 trials because the female legs appeared to be injured. 1 trial was removed because the couple already started copulating at recording start. 14 trials were removed because video-recording got unintentionally interrupted before the male adopted a stable copulation posture on top of the female. In these trials it was uncertain if the couple would adopt a stable copulation posture or not. Other movies, where video-recording got unintentionally interrupted were analyzed when the male had reached a stable copulation posture. The movie ends are labelled (yellow) in the ethogram (Figure S7). In total, we observed stable copulation in 34 / 63 trials. One trial was excluded from assessment of the copulation posture because the couple mated being attached to the inner side of the transparent plastic cover and was only observed in ventral view so that the male head and the female scutellum were not visible (see below). In order to perform a randomized mating position analysis with respect to the male genotype, movie names were replaced by a seven-digit random number (supplementary dataset S8). Images were automatically extracted from each movie at 10 sec after the male had reached a stable copulation posture and then every 5 min until copulation or acquisition ended. Image extraction from movies was prepared according to (Acurio et al., 2019) using custom R and bash scripts and the avconv software (libav tools, https://www.libav.org). Position analysis was also carried out according to (Acurio et al., 2019). We positioned three landmarks on the female and male body: P1 was the anterior medial tip of the female head (Figure 4, Figure S8, red point), P2 the distal tip of the female scutellum (Figure 4, Figure S8, blue point) and P3 the most posterior medial point of the male head (Figure 4, Figure S8, open black circle). Landmarks were placed manually on each image using imageJ (supplementary dataset 8). Data analysis was done with a custom R script to rotate landmark positions and to scale landmark distances, so that all P2 points were super-imposed in the diagram origin and all P1 points were aligned to position (0,1)

(Figure S8 B-G). The angle (α) between lines (P1-P2) and (P2-P3) (Figure 4, Figure S8) was used to measure sidedness of the mating position on each extracted image, positive values indicating right-sidedness and negative values left-sidedness. Repeatability of landmark positioning was assessed by two independent rounds of coordinate acquisition (Figure S8). Variation in angle estimates was found to be attributable mostly to individual images and not to replicate measurement (Figure S8A). Hypothesis testing was performed in R to evaluate the overall sidedness of mating posture among groups of trials with different male genotypes (Figure 4), with the null hypothesis: angle = 0, using the functions glm for generalized linear model fits, and the function glht() to derive estimated contrasts ($myo1D^{wt/wt}$ estimate = 16.297, standard error = 2.417, z = 6.742, $p < 4.7 \times 10^{-11}$, $myo1D^{wt/mut}$ estimate = 17.912, standard error = 2.640, z = 6.784, $p < 3.52 \times 10^{-11}$, $myo1D^{mut/mut}$ estimate = 39.627, standard error = 2.764, z = 14.335, $p < 2 \times 10^{-16}$).

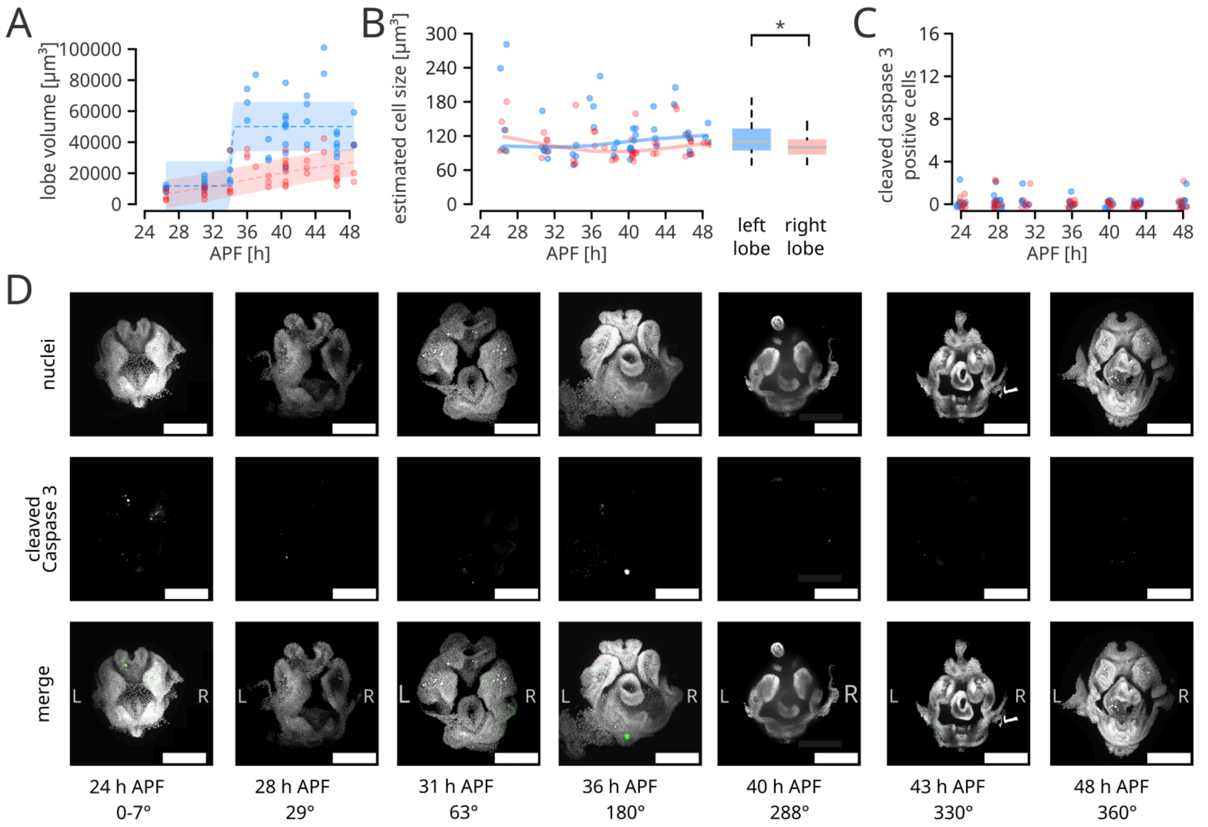


**Figure S1: Mean lobe volume, lobe cell size and cell death. (A)** Estimated lobe volume (left lobe in light blue and right lobe in pink) from contour analysis of confocal fluorescence microscopy scans of developing genitalia at different time points after puparium formation. 4-parameter logistic regressions and standard errors are shown as dashed lines and colored shades in light blue and red, respectively. **(B)** Estimated cell size of left lobe (light blue) and right lobe (pink) cells at different timepoints after puparium formation, calculated as the ratio of lobe volume and the number of nuclei counted therein. The trendline is a lowess regression curve. The sample distribution is summarized as boxplots on the right for all time points. The estimated left lobe cell size in the left lobe (light blue) is slightly larger than in the right lobe (t-test, $t = 2.2973$, $df = 75.736$, $p = 0.02436$), as indicated by the star and the bracket. **(C)** Number of cleaved caspase 3 cells in the left lobe (light blue) and right lobe (pink) during pupal development. **(D)** Fixed *D. pachea* male genitalia at different time-points after puparium formation. Nuclei were stained with DAPI (grey, first and third rows) and apoptotic cells with an anti-cleaved Caspase 3 antibody (white in second and green in third row). Left and right are indicated with L and R, respectively. The scale is 100 µm.

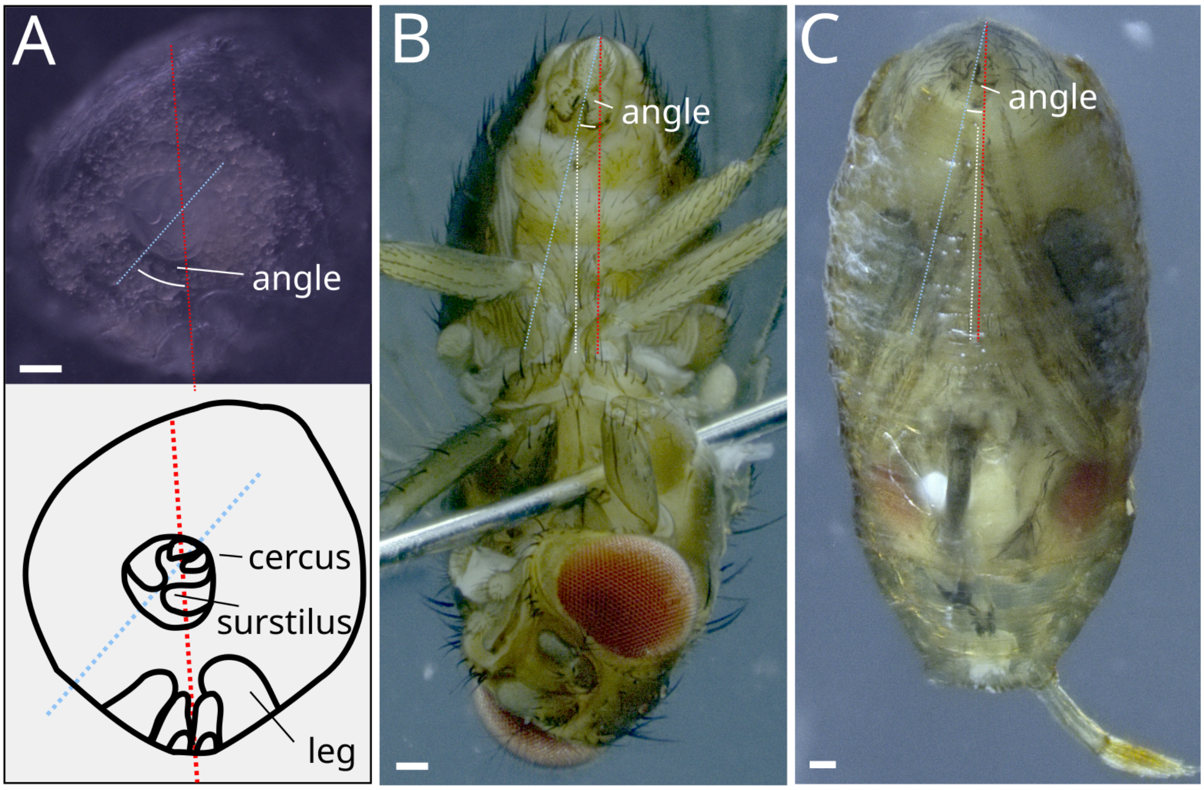


**Figure S2: Quantification of genitalia orientation. (A)** Male genitalia orientation approximation on dissected posterior parts of male pupa for data in Figure 1E. Genitalia on dissected pupae are outlined. Orientation was measured as the angle between the dorso-ventral axis of male genitalia (blue dashed line) estimated as the midline between the paired cerci and surstili, and the male dorso-ventral body axis (red dashed line) approximated as the medial line of the pupa relative to the ventrally located legs. **(B)** Genitalia orientation in adult males was measured as the angle between the dorsoventral axis of genitalia (blue dashed line) and the antero-posterior axis of the abdomen (white and red dashed lines)relative to the ventrally located legs. **(C)** Genitalia orientation in pharate adult pupae were measured as in (B). The scale is 100 µm.

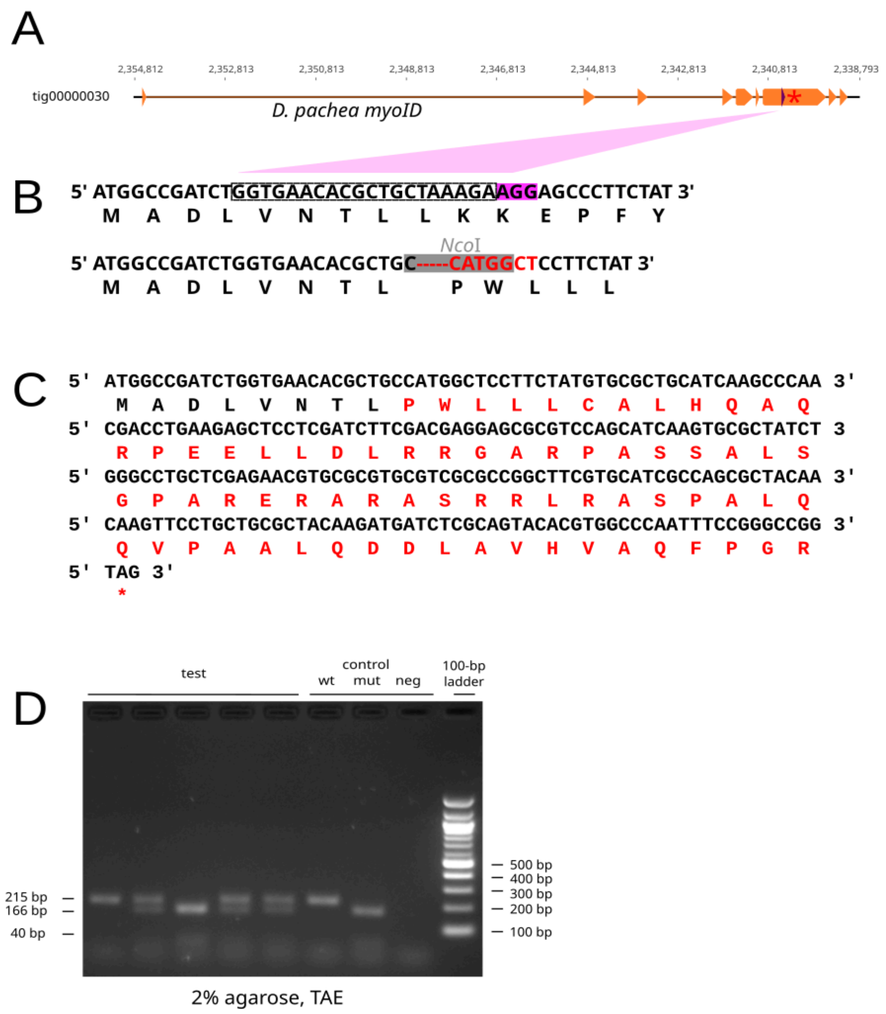


**Figure S3: Identification of the *myo1D* mutation. (A)** Annotation of the *myo1D* coding sequence (orange). Orange arrows indicate coding exons, numbers indicate the nucleotide positions relative to contig tig00000030 of the *D. pachea* genome assembly (Bioproject PRJNA611543), the violet arrow indicates the CRISPR target site and the red star indicates the location of the premature stop codon in the *myo1D* mutant, adapted from Geneious 11.1.5 (https://www.geneious.com). **(B)** Nucleotide and amino acid sequences at the CRISPR target site. The upper line corresponds to the *D. pachea* wild-type *myo1D* sequence, the lower line to the CRISPR/Cas-9 induced frameshift mutation. Outlined framed positions indicate the single guide RNA (sgRNA) sequence to target *myo1D*, dark-violet highlights the protospacer adjacent motif (PAM). The mutated positions are in red, the *Nco*I endonuclease motif is highlighted in grey. Single letter amino acid abbreviations are presented beneath each codon. **(C)** The mutation causes a frameshift. Altered amino acid sequence is indicated by red single letter amino acid abbreviations. **(D)** Gel electrophoresis of *Nco*I digested PCR products, used to distinguish *D. pachea* male genotypes. Test indicates lanes corresponding to genomic DNA of *D. pachea* *myo1D*$^{wt/wt}$, *myo1D*$^{wt/mut}$, *myo1D*$^{mut/mut}$ males males. Controls *myo1D*$^{wt/wt}$ (wt), *myo1D*$^{mut/mut}$ (mut) males and a negative control amplified from water (neg). The 100 bp ladder is presented on the right. Relevant DNA fragment sizes are indicated at the sides of the image, the agarose concentration and gel electrophoresis buffer is annotated below the image.

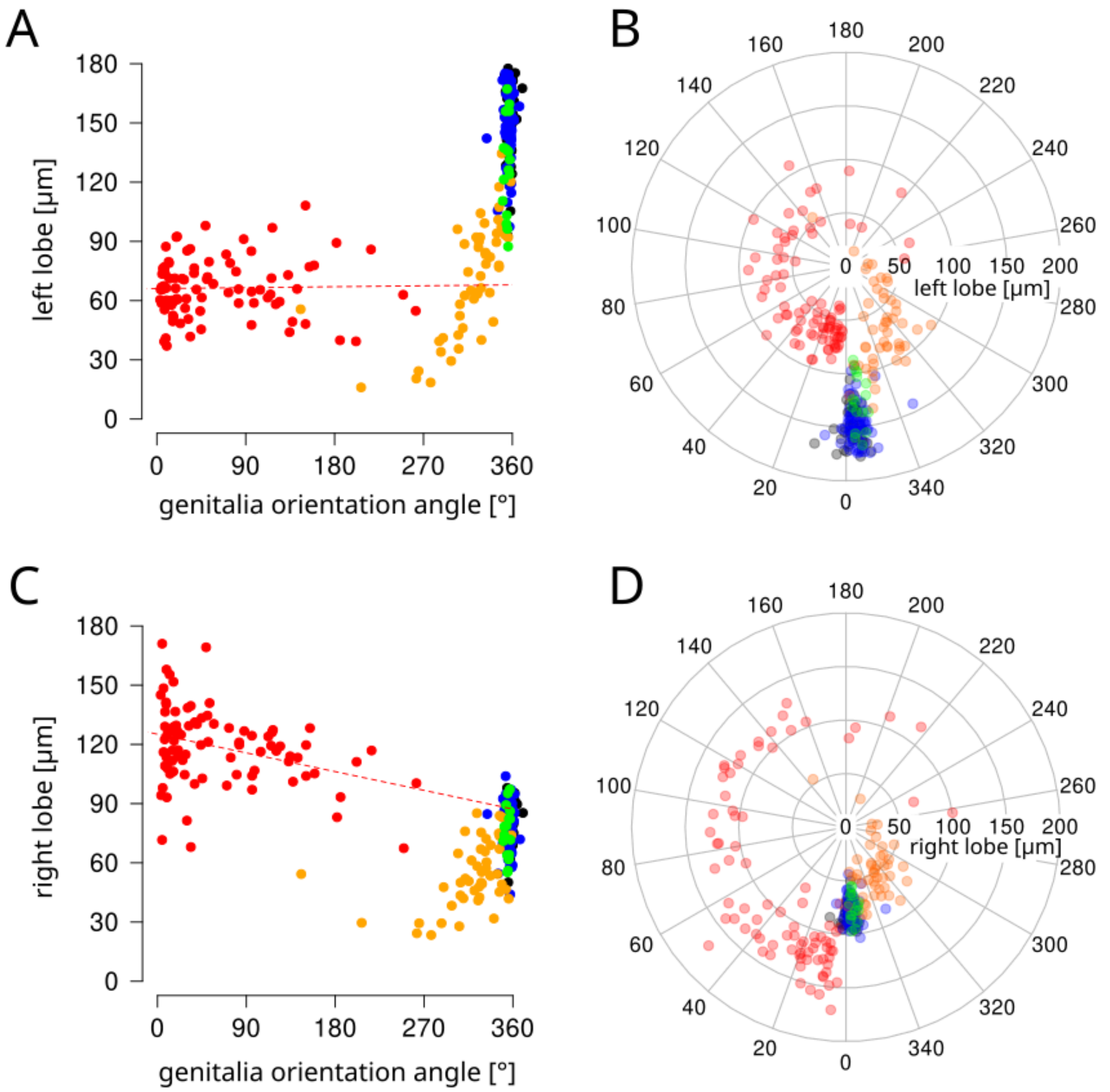


**Figure S4: Distribution of lobe lengths at different genitalia orientation angles,** of *myo*$^{wt/wt}$ males (black), *myo*$^{wt/mut}$ males (blue), and *myo*$^{mut/mut}$ males (red), pyriproxyfen treated individuals (orange) and solvent control individuals (green). **(A)** Left lobe length on a scatter plot. The dashed red line is a linear regression including data from *myo*$^{mut/mut}$ males (Pearson correlation, coefficient=0.022, t= 0.201, df=84 p=0.841), **(B)** left lobe length on a polar plot, **(C)** right lobe lengths on a scatter plot. The dashed red line is a linear regression including data from *myo*$^{mut/mut}$ males (Pearson correlation coefficient=-0.338, t=-3.286, df=84, p=0.0015). **(D)** right lobe length on a polar plot.

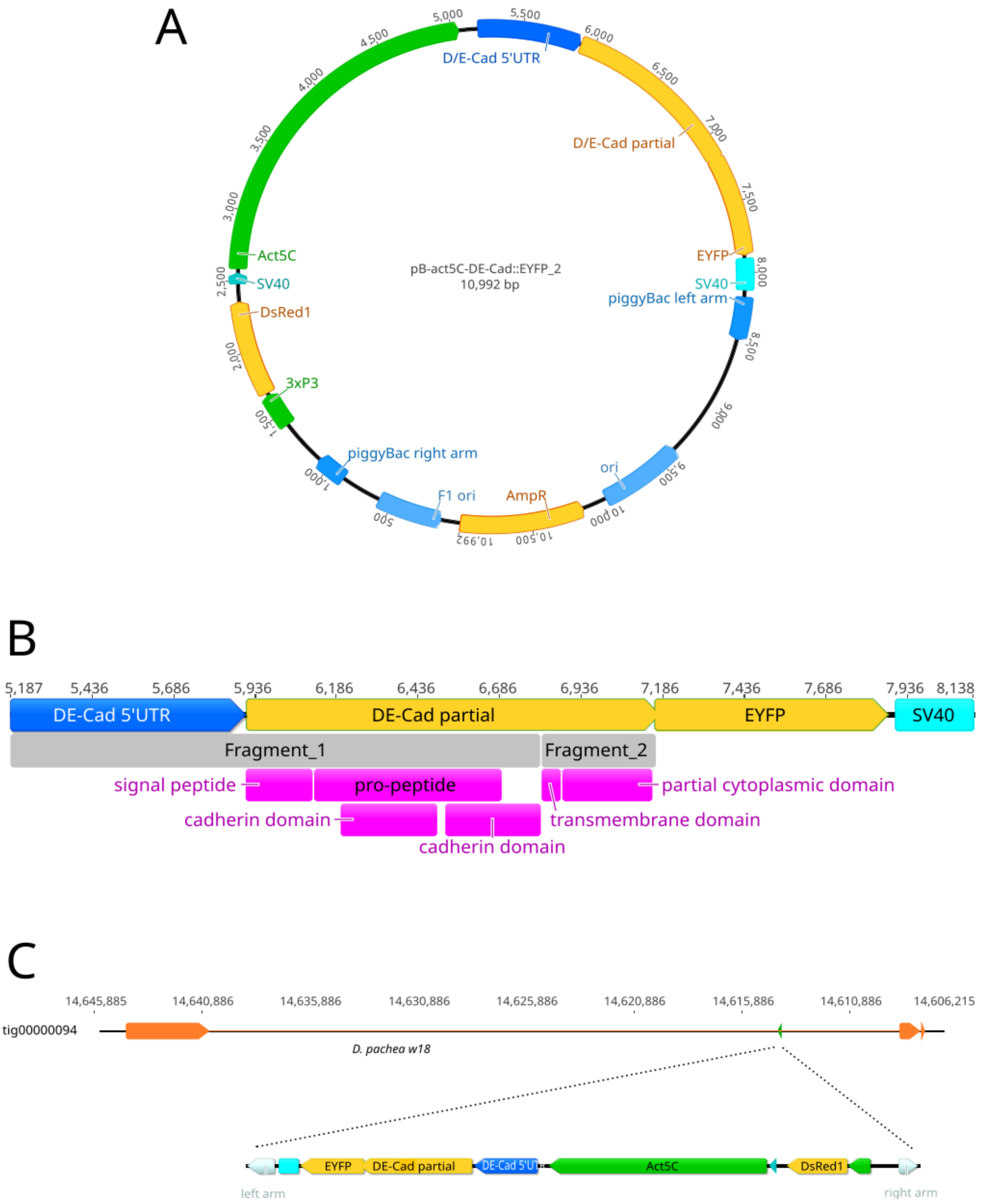


**Figure S5: DE-cadherin-EYFP membrane marker. (A)** Plasmid map of construct act5C::DE-Cad-EYFP_2. **(B)** Diagram of the plasmid region that encodes the partial D/E cadherin (*shg*) coding sequence, fused to the coding sequence of EYFP. Grey blocks labeled Fragment_1 and Fragment_2 correspond to partial *D. melanogaster shg* coding portions, pink blocks indicate encoded DE-cadherin protein domains. **(C)** Localization of the act5C::DE-Cad-EYFP_2 insert inside contig tig00000094. The coding exons of the *D. pachea wheeler 18* locus is indicated by orange arrows. The insertion site of act5C::DE-Cad-EYFP_2 is labeled with a green arrow. Drawings were adapted from Geneious 11.1.5 (https://www.geneious.com).

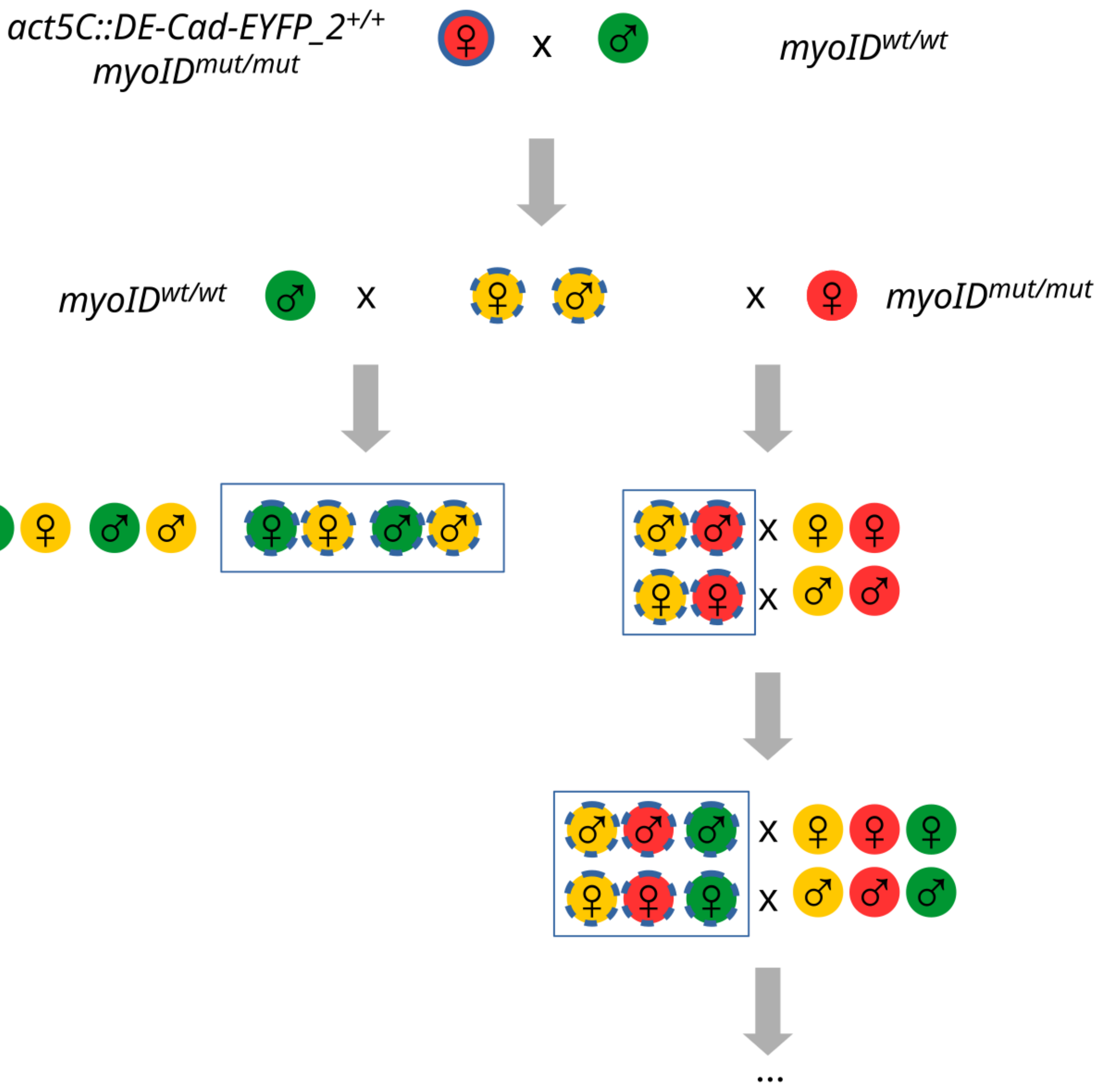


**Figure S6: Crossing scheme for time-lapse microscopy samples.** Homozygous females [*myo1D*$^{mut/mut}$, act5C::DE-Cad-EYFP_2$^{+/+}$] were crossed to wild-type males [*myo1D*$^{wt/wt}$, act5C::DE-Cad-EYFP_2$^{-/-}$]. The female F1 progeny was crossed back to wild-type males and male progeny was crossed to *myo1D*$^{mut/mut}$ mutant males [*myo1D*$^{mut/mut}$, act5C::DE-Cad-EYFP_2$^{-/-}$]. The progeny of the latter cross was then maintained by sibling crosses: females [act5C::DE-Cad-EYFP_2$^{+/-}$] with males [act5C::DE-Cad-EYFP_2$^{-/-}$] and females [act5C::DE-Cad-EYFP_2$^{-/-}$] with males [act5C::DE-Cad-EYFP_2$^{+/-}$]. The presence of the act5C::DE-Cad-EYFP_2 insertion was visible under a fluorescence stereomicroscope by the presence of red fluorescence in the adult eyes. Colored circles indicate *myo1D* genotypes, green: myo1D$^{wt/wt}$, yellow: *myo1D*$^{wt/mut}$, red: *myo1D*$^{mut/mut}$. Outlines indicate act5C::DE-Cad-EYFP_2 insertion, absent: no insertion, dashed: heterozygous insertion, full: homozygous for the insertion. Open squares indicates genotypes of individuals used for time-lapse imaging. Times-lapse imaging was performed on individuals originating from the first three successive generations (rotprob1, rotprob2, rotprob3 in supplementary dataset 5).

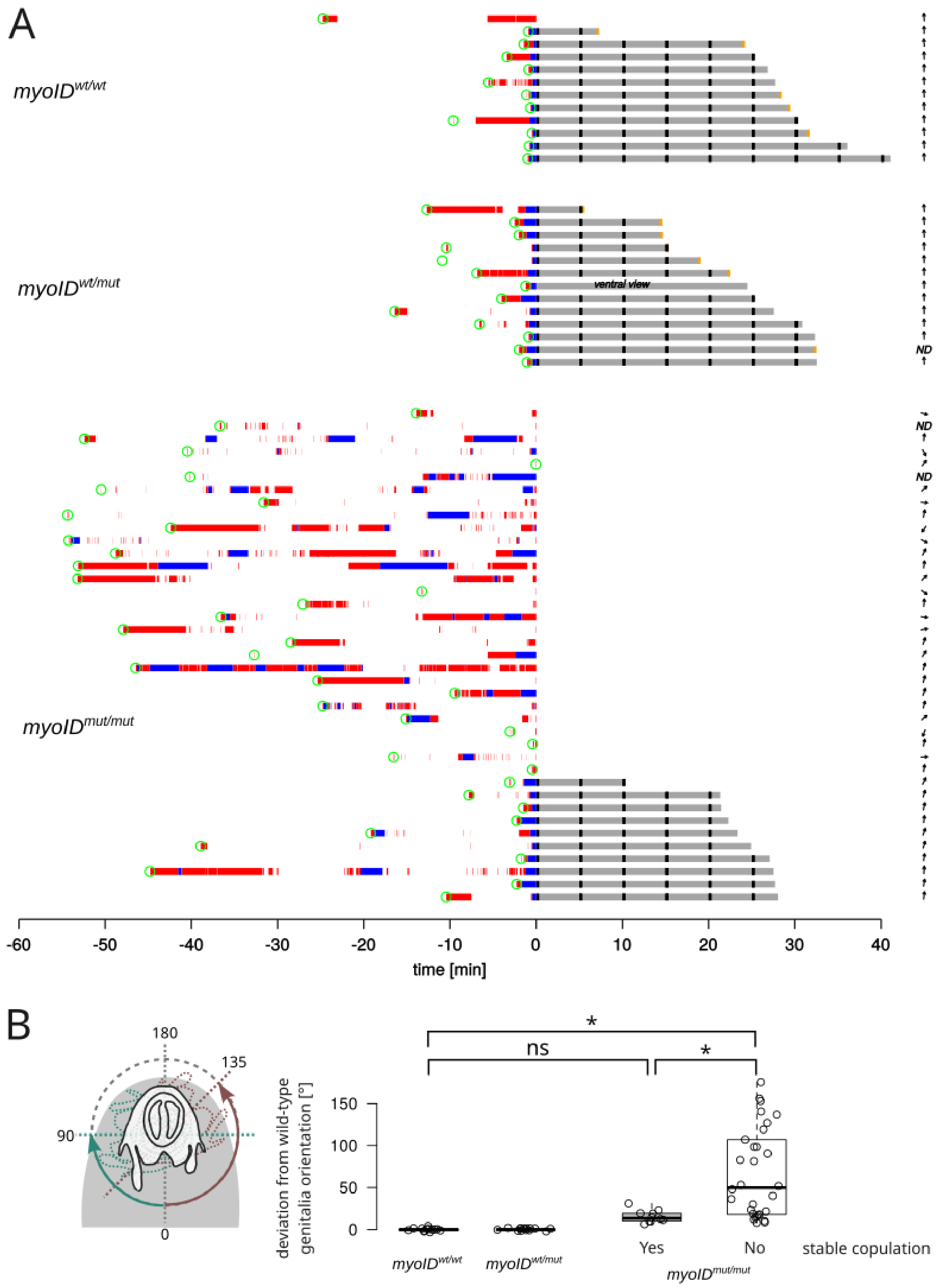


**Figure S7: Mating behavior of *D. pachea* males differing in *myo1D* genotype. (A)** Ethogram, (Supplementary datafile 7). Each line presents a mating trial. Green circles is courtship start, red segments is male licking behavior, blue segments are mounting attempts, grey segments represent stable copulation, black tick marks indicate mating position measurement (supplementary datafile 8). Copulation was observed in "ventral view" in one trial. Yellow labels show movie-ends before copulation had ended. Arrows indicate male genitalia orientation (supplementary datafile 4). ND stands for "not determined". **(B)** Male genitalia absolute orientation differences relative to mean *myo1D*$^{wt/wt}$ orientation. Standardized values range from 0 - 180°. The drawing on the left illustrates male genitalia orientations on the male ventral abdomen (grey shade) with two deviation calculation examples in green and red, for angle deviation to the left and right, respectively. No significant orientation differences were found between *myo1D*$^{wt/wt}$ males, *myo1D*$^{wt/mut}$ males and *myo1D*$^{mut/mut}$ males that achieved stable copulation, but orientation significantly deviated in *myo1D*$^{mut/mut}$ males that did not achieve stable copulation. Significance is indicated as ns for "not significant' and * for $p < 0.001$ or lower, and was tested with Bonferroni corrected pairwise contrasts on a generalized linear model fit (orientation ~ group) for the hypothesis of equal orientation (*myo1D*$^{wt/wt}$, *myo1D*$^{mut/mut}$ [stable copulation]): estimate = 15.6113, standard error = 15.2179, z-value = 1.026, p = 1; (*myo1D*$^{wt/wt}$, *myo1D*$^{mut/mut}$ [no stable copulation]): estimate = 66.1209, standard error = 12.0133, z-value = 5.527, $p < 0.00001$, (*myo1D*$^{mut/mut}$, *myo1D*$^{mut/mut}$ [stable copulation]): estimate = 50.7909, standard error = 13.2108, z-value = 3.845, $p < 0.001$).

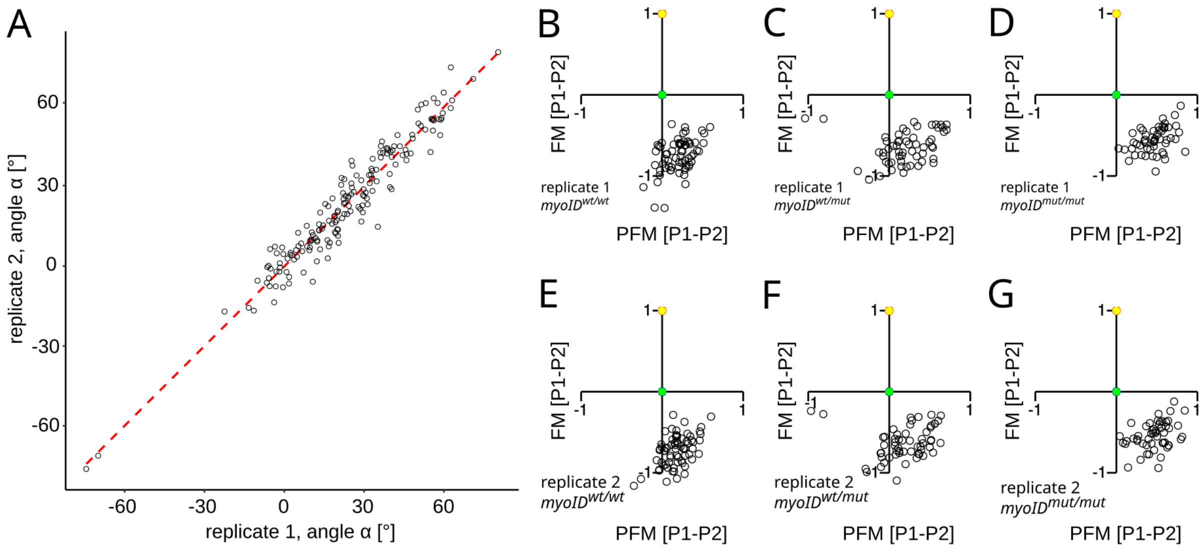


**Figure S8: Quantification of copulation postures. (A)** Two sets of mating angle measurements (Figure 4A), done by two persons on a randomized image dataset, to compare repeatability of landmark positioning. Data analyses are highly correlated (Pearson correlation: coefficient = 0.967, t = 51.773, df = 206, p < 2.2 x 10-16, r = 0.967). The red dashed line is a linear regression. Variation in angle estimates was found to be attributable mostly to individual images of the different movies (parameter: image) but not to replicate measurements (parameter: replicate) (ANOVA, angle ~ image + replicate, image: df1 = 176, df2 = 1, image: F = 58.74, $p < 10^{-16}$, replicate: F = 0.148, p = 0.701). **(B-G)** Visualization of landmark positions (P1: yellow points, P2: green points, P3: open circles, Figure 4A) of couples with *myo1D*$^{wt/wt}$, *myo1D*$^{wt/mut}$, *myo1D*$^{mut/mut}$ males of each replicate analysis. Landmark positions are rotated and scaled relative to the distance of P1 and P2 on the female dorsum. The FM axis corresponds to the female midline, PFM axis describes the perpendicular axis to the female midline.

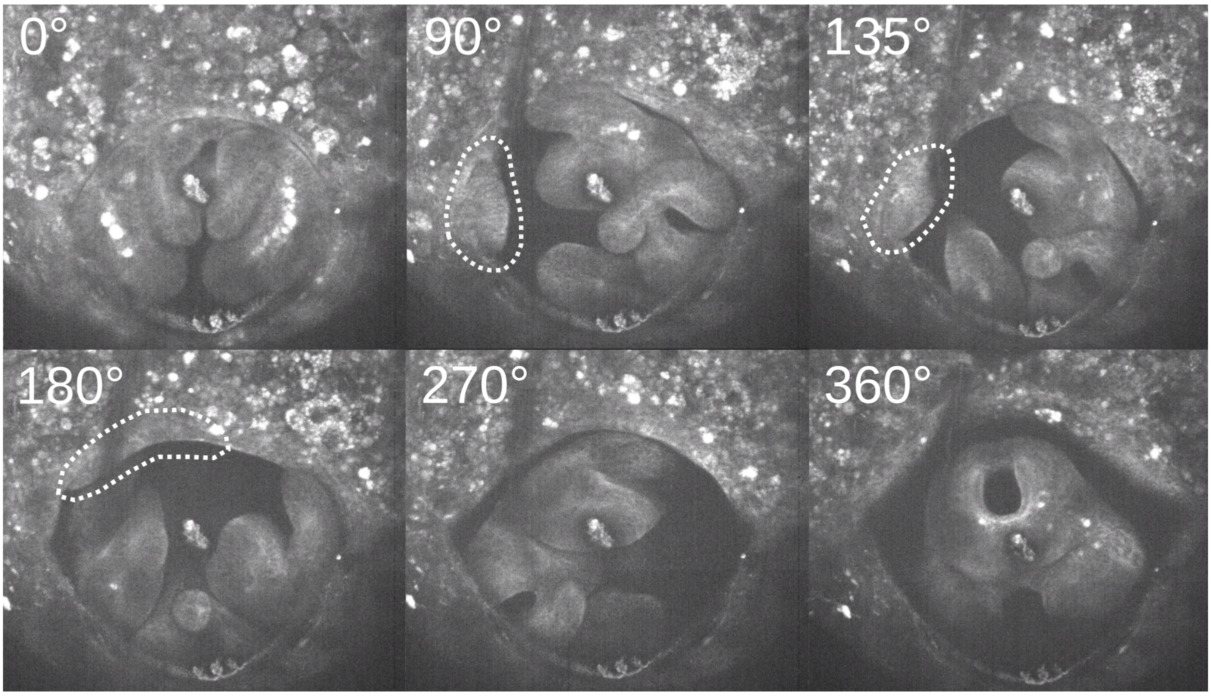


**Figure S9: Genitalia rotation involves surrounding tissue agglomeration.** Snapshots from movie 1, time-lapse microscopy of a *D. pachea* heterozygous *myo1D*$^{wt/mut}$ male (supplementary dataset 4, individual 331_dev). The rotation angle relative to the orientation at rotation start is indicated on each panel. Tissue agglomeration is outlined with a dashed white line.

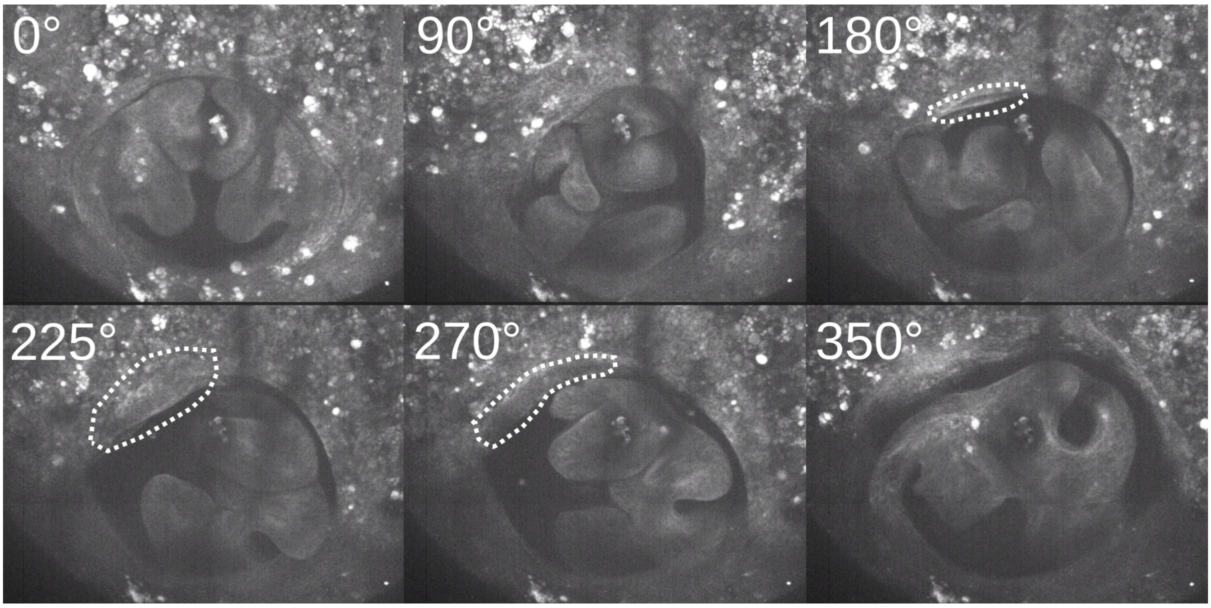


**Figure S10: Tissue agglomeration during genitalia rotation in a *D. pachea myo1D*$^{mut/mut}$ male.** Snapshots from time-lapse microscopy, movies 2 (Supplementary dataset 4, individual 345_dev). The rotation angle relative to the orientation at rotation start is indicated on each panel. Tissue agglomeration is outlined with a dashed white line.

**Table S1. Oligonucleotides for PCR and preparation of CRISPR sgRNA.** Tan °C corresponds to annealing temperatures of thermocycle programs used in our experiments.

| oligo-nucleotide | sequence | Tan [°C] | reference |
|---|---|---|---|
| pacMyo1D-retro | CCTAATATGATACACCGAACTAT | 50 | this study |
| pacMyo1D-debut | GAGCAAAACAAAAACGCAGAAACAT | 60 | this study |
| pacMyo1D-stop | CGCTCAGCAGCTATCAGCTCTAA | 60 | this study |
| 2F_Dpac_m31DF | TCGCCAGGACTTCCGCATTA | 60 | this study |
| 4F_Dpac_m31DF | CCAATTCGCAGGACAACAG | 60 | this study |
| 1R_Dpac_m31DF | GCCGATTACCGAGTTGAATCT | 60 | this study |
| Dpa2Rbis3 | CGCAGCAGGAACTTGTTGTA | 60 | this study |
| Dpa_myo1DCRISPR_F | GAAATTAATACGACTCACTATAGGTGAACACGCTGCTAAAGAGTTTTAGAGCTAGAAATAGC | 60 | this study |
| CRISPR-sgR | AAAAGCACCGACTCGGTGCCACTTTTTCAAGTTGATAACGGACTAGCCTTATTTTAACTTGCTATTTCTAGCTCTAAAAC | 60 | (Bassett et al., 2013) |
| 1F_Dpac_m31DF | CGCCAACTGGACCTGCATAA | 60 | this study |
| Myo1D_3F-genoF1 | ATTATCCAGCGGTACATGGCCGAT | 70-60 (touch-down) | this study |
| m_DECad_R1 | CGGCTGGCGAAGATTCCTA | 62 | this study |
| m_DECad_F1 | CCTGCAGGTCGACCACTCGAGTGTTGCATTGTGTTTTTGC | 62 | this study |
| mDECad_R2 | CAATGATGAAAAAATGATGGCGGCTTAATTG | 62 | this study |
| mDECad_F2 | GCCATCATTTTTCATCATTGCGATCATCGTATG | 64 | this study |
| m_DECad_R3 | TTGGGAGCTCTCCCATACTCGAGCACATCGTCCACGGTTGTG | 64 | this study |
| act5C_F1 | GGTTACGGCACTAGAGCGGCCGCGCAATTCTATATTCTAAAAACACAAATGATACTTCTAAAA | 60 | this study |
| act5C_R | AAAACATACATATGTACATGTGTTTATAGTGGGTTTATTAG | 60 | this study |
| act5C_F2 | TATAAACACATGTACATATGTATGTTTTGGCATACAATGAGTAGTTG | 65 | this study |
| SV40_R | TACCGTCGACCTCGAGAGATCTGATCCAGACATGATAAGATACATTGATGAGTTTGGACAAACCACAACTAGAA | 65 | this study |
| InvPCR_F1 | CGGCGACTGAGATGTCCTAAAT | 64 | this study |
| InvPCR_R1 | GCGGTAAGTGTCACTGATTTTGAACTATAA | 64 | this study |
| InvPCR_F2 | CGCGCTATTTAGAAAGAGAGAGCAATATTT | 64 | this study |
| InvPCR_R2 | GACCGCGTGAGTCAAAATGA | 64 | this study |

**Legends for Movies 1 to 2 and Datasets S1 to S8**

**Supplementary Dataset 1.** Confocal microscopy analysis of fixed male genitalia during metamorphosis.

**Supplementary Dataset 2.** Genitalia rotation angles of male genitalia during metamorphosis.

**Supplementary Dataset 3.** Viability of D. pachea and D. melanogaster to pyriproxyfen application.

**Supplementary Dataset 4.** Male *D. pachea* genitalia morphology and orientation, spreadsheet with length and orientation measurements of genital morphology of male *D. pachea* flies.

**Supplementary Dataset 5.** Time-lapse microscopy trials, summary of trials and conditions for time-lapse microscopy.

**Supplementary Dataset 6.** Single couple mating trials, summary of *D. pachea* single couple mating trials.

**Supplementary Dataset 7.** Courtship and copulation behavior, annotated courtship and copulation behavior. The data is summarized in Fig. S6.

**Supplementary Dataset 8.** Copulation position coordinates, position landmarks to compare copulation postures. The data is the main input file for Fig. 2.

**Movie 1.** Clockwise genitalia rotation, acquired by time-lapse microscopy of a *D. pachea* heterozygous $myo1D^{wt/mut}$ mutant male.

**Movie 2.** Counter-clockwise genitalia rotation, acquired by time-lapse microscopy of a *D. pachea* homozoygous $myo1D^{mut/mut}$ mutant male.

### SI References


Acurio, A.E., Rhebergen, F.T., Paulus, S., Courtier-Orgogozo, V., Lang, M., 2019. Repeated evolution of asymmetric genitalia and right-sided mating behavior in the Drosophila nannoptera species group. BMC Evolutionary Biology 19, 109. https://doi.org/10.1186/s12862-019-1434-z

Bainbridge, S.P., Bownes, M., 1981. Staging the metamorphosis of Drosophila melanogaster. Development 66, 57–80.

Bassett, A.R., Tibbit, C., Ponting, C.P., Liu, J.-L., 2013. Highly efficient targeted mutagenesis of Drosophila with the CRISPR/Cas9 system. Cell reports 4, 220–228.

Gibson, D.G., Young, L., Chuang, R.-Y., Venter, J.C., Hutchison, C.A., Smith, H.O., 2009. Enzymatic assembly of DNA molecules up to several hundred kilobases. Nature methods 6, 343–345.

Heed, W.B., Kircher, H.W., 1965. Unique sterol in the ecology and nutrition of Drosophila pachea. Science 149, 758–761.

Horn, C., Offen, N., Nystedt, S., Häcker, U., Wimmer, E.A., 2003. piggyBac-based insertional mutagenesis and enhancer detection as a tool for functional insect genomics. Genetics 163, 647–661.

Langmead, B., Trapnell, C., Pop, M., Salzberg, S.L., 2009. Ultrafast and memory-efficient alignment of short DNA sequences to the human genome. Genome Biol 10, R25. https://doi.org/10.1186/gb-2009-10-3-r25

Lefèvre, B.M., Catté, D., Courtier-Orgogozo, V., Lang, M., 2021. Male genital lobe morphology affects the chance to copulate in Drosophila pachea. BMC ecology and evolution 21, 1–13.

Matz, M.V., Fradkov, A.F., Labas, Y.A., Savitsky, A.P., Zaraisky, A.G., Markelov, M.L., Lukyanov, S.A., 1999. Fluorescent proteins from nonbioluminescent Anthozoa species. Nature biotechnology 17, 969–973.

McAlpine, J.F., 1981. Morphology and terminology-adults. Manual of nearctic diptera 1, 9–63.

Nagy, O., Nuez, I., Savisaar, R., Peluffo, A.E., Yassin, A., Lang, M., Stern, D.L., Matute, D.R., David, J.R., Courtier-Orgogozo, V., 2018. Correlated evolution of two copulatory organs via a single cis-regulatory nucleotide change. Current Biology 28, 3450–3457.

Ochman, H., Gerber, A.S., Hartl, D.L., 1988. Genetic applications of an inverse polymerase chain reaction. Genetics 120, 621–623.

Oda, H., Uemura, T., Harada, Y., Iwai, Y., Takeichi, M., 1994. A Drosophila homolog of cadherin associated with armadillo and essential for embryonic cell-cell adhesion. Developmental biology 165, 716–726.

Pacquelet, A., Lin, L., Rørth, P., 2003. Binding site for p120/δ-catenin is not required for Drosophila E-cadherin function in vivo. Journal of Cell Biology 160, 313–319. https://doi.org/10.1083/jcb.200207160

Pitnick, S., 1993. Operational sex ratios and sperm limitation in populations of Drosophila pachea. Behav Ecol Sociobiol 33. https://doi.org/10.1007/BF00170253

Spéder, P., Ádám, G., Noselli, S., 2006. Type ID unconventional myosin controls left–right asymmetry in Drosophila. Nature 440, 803–807.

Spieth, H.T., 1952. Mating behavior within the genus Drosophila (Diptera). American Museum of Natural History.

Suvorov, A., Kim, B.Y., Wang, J., Armstrong, E.E., Peede, D., D'agostino, E.R., Price, D.K., Waddell, P.J., Lang, M., Courtier-Orgogozo, V., 2022. Widespread introgression across a phylogeny of 155 Drosophila genomes. Current Biology 32, 111–123.

Thummel, C.S., Boulet, A.M., Lipshitz, H.D., 1988. Vectors for Drosophila P-element-mediated transformation and tissue culture transfection. Gene 74, 445–456.
